# Oxalic acid binds to gustatory receptor Gr23a and inhibits feeding in the brown planthopper

**DOI:** 10.1101/2021.10.14.464394

**Authors:** Kui Kang, Mengyi Zhang, Lei Yue, Weiwen Chen, Yangshuo Dai, Kai Lin, Kai Liu, Jun Lv, Zhanwen Guan, Shi Xiao, Wenqing Zhang

**Affiliations:** State Key Laboratory of Biocontrol and School of Life Sciences, Sun Yat-sen University, Guangzhou 510275, Guangdong, China; College of Biology and Agriculture, Zunyi Normal University, Zunyi 563006, Guizhou, China

**Author notes:** These authors contributed equally to this work (Kui Kang, Mengyi Zhang).

**Keywords:** ligand identification, gustatory receptor, antifeedant, aversive behaviour, oxalic acid

## Abstract

Plants produce diverse secondary compounds as natural protection against microbial and insect attack. Most of these compounds, including bitters and acids, are sensed by insect gustatory receptors (Grs). Acids are potentially toxic to insects, but there are few reports on sour compounds as ligands of insect Grs. Here, using two different heterologous expression systems, the insect Sf9 cell line and the mammalian HEK293T cell line, we started from crude extracts of rice (*Oryza sativa*) and successfully identified oxalic acid (OA) as a ligand of NlGr23a, a Gr in the brown planthopper *Nilaparvata lugens*. The antifeedant effect of OA on the brown planthopper was dose dependent, and *NlGr23a* is essential for OA’s antifeedant activity in both artificial diets and rice plants. *NlGr23a* is also indispensable for tarsal OA sensing. To our knowledge, OA is the first identified ligand starting from plant crude extracts and the first known strong acid for insect Grs. These findings on rice-planthopper interactions will be of broad interest for pest control in agriculture and also for better understanding of how insects select host plants.

**Research organism:** *Nilaparvata lugens*

## Introduction

Plant visual and chemical cues are important for host location and host acceptance of phytophagous insects. The cabbage root fly, *Delia radicum*, makes use of leaf colors to discriminate among host plants (Prokopy *et al*., 1983). Host selection in oligophagous species is involved with the balance of phagostimulatory and deterrent inputs with the addition of a specific chemical sign stimulus (Chapman, 2003). Understanding the mechanisms underlying this selection strategy is of practical importance in agriculture for pest control. For instance, tandem deployment of the napier grass, which attracts more oviposition by stemborer moths than maize but not suitable for survival, and the molasses grass, which causes over 80% reduction in stemborer infestation of maize, can interfere host selection by stemborers (Khan *et al.*, 2014). The combination of stimuli that have negative effects on food selection by pests and positive effects on diverting pests from the protected resource to a trap can reduce pest abundance on the protected resource, which is called the push-pull strategy (Cook *et al.*, 2007). Plant-derived antifeedants can be used as push components of the push-pull strategy (Cook et al., 2007). Identification of receptors sensing these chemicals will definitely improve our understanding of how insects select host plants.

To date, most identified antifeedants are secondary metabolites, which are toxic or deterrent to insects (Yoshihara *et al.*, 1980; Chapman, 2013; Douglas, 2017). As an example, oxalic acid (OA), the simplest dicarboxylic acid common in plant tissues, provides protection for plants against insects and foraging animals, including the brown planthopper (BPH) and aphids (Massonie, 1980; Yoshihara *et al.*, 1980; Karatolos and Hatcher, 2008). Some antifeedants (e.g. tricin) could be even used as indicators of crop resistance to target insect pests, and are therefore potentially valuable for breeding resistant crop varieties (Zhang *et al.*, 2015).

In insects, gustatory receptor neurons (GRNs), mainly found in gustatory sensilla, are able to convert the chemical signal into an electrical one, and transmit it to higher-order brain structures for processing, which in turn dictates behaviour (Scott, 2018). Secondary compounds as deterrents stimulate a subset of bitter GRNs, which then inhibit feeding activity or induce repulsion (Chapman, 2003). Different secondary metabolites elicited different responding patterns of GRNs in the two *Helicoverpa* species might mediate the dietary acceptance and host range (Sun *et al.*, 2021).With the decoding and publication of the genomes of *Drosophila melanogaster* and other insect species, fruitful molecular studies on insect gustation have been enabled. Gustatory receptors (Grs), expressed in GRNs, are required for responses to specific tastants. Grs mainly include sweet, bitter and CO_2_ receptors (Liman *et al.*, 2014). Although diverse sugars and bitter compounds have been identified as ligands of insect Grs (Chen and Dahanukar, 2020), many Grs have yet to be functionally annotated, particularly the Grs sense acids. DmGr64e in *D. melanogaster* was reported to sense fatty acids such as hexanoic acid and octanoic acid (Kim *et al.*, 2018). However, there are currently no reports of strong acid as ligands of insect Grs.

The BPH *Nilaparvata lugens*, causing extensive damage by sap feeding and transmitting viruses, feeds solely on cultivated rice and its allied wild forms such as *Oryza perennis* and *O. spontanea* (Sogawa, 1982). Its gustatory sensilla are mainly located in the small passageway leading from the food duct to the cibarium and the stylet groove on the labial tip, and probably regulate the initial stages of probing and in the maintenance of sap ingestion (Foster *et al.*, 1983a; Foster *et al.*, 1983b). In our previous study, 32 of *N. lugens* Gr sequences (*NlGrs*) were obtained following analyses of the BPH genome and transcriptomes (Kang *et al.*, 2018). Here, we selected a functionally unknown gene, *NlGr23a*, which is related to BPH fecundity. Through a multi-stage bioassay-directed fractionation of rice crude extracts, we identified OA as a ligand of NlGr23a. We also found that *NlGr23a* is essential to the antifeedant activity of OA in both artificial diets and rice plants and the modulation of the tarsal repulsion for OA. To our knowledge, OA is the first identified ligand starting from plant crude extracts, as well as the first known strong acidic ligand for insect Grs.

## Results

### Oxalic acid is a ligand of NlGr23a

To identify the specific ligand of NlGr23a, we cloned the full-length *NlGr23a* open reading frame (GenBank accession No. MT387198), which encodes 451 amino acid residues with seven predicted transmembrane domains (see Appendix 1—figure 1A). Then NlGr23a-expressing stable *Spodoptera frugiperda* (Sf9) cell line was established by antibiotic selection for at least 10 passages. Expression of the stably transfected gene was verified by immunoblotting and RT-PCR (see Appendix 1—figures 1B and C).

We carried out 4 rounds of ligand screening to search for potential ligands of NlGr23a from crude extracts of rice stems and leaves. The initial calcium imaging results revealed that the ethyl acetate (EAC) and methyl alcohol (MeOH) fractions caused significantly higher calcium ion release/concentration (i.e. cellular response) as indicated by the ratio of cytoplasmic fluorescence intensities (Figure 1A). The EAC fraction was then further fractionated into three portions for the next round of screening; Sf9 cells expressing *NlGr23a* showed a response only to fraction 2 (time of retention, *t_R_ =* 20–35 min, Figure 1B and see also Appendix 1—figure 2A for HPLC chromatogram of the EAC fraction). This fraction was subjected to further fractionation with acetonitrile (ACN) and we found that only the subfraction collected in 5–10 min specifically evoked Ca^2+^ release (Figure 1C and see also Appendix 1—figure 2B for HPLC chromatogram of the subfraction). The bioassay-positive subfraction was isolated and analysed using gas chromatography-mass spectrometry (GC-MS). Four compounds, oxalic acid (OA), glycerol, phthalic acid, and trisiloxane, were identified (Figure 1D and see also Appendix 1—figure 2C for GC-MS chromatogram of the control solution). As the peak intensity of phthalic acid was quite weak, and trisiloxane was probably an artefact of culture vessel silanization (see Appendix 1—table 1 for the GC-MS analysis details), OA and glycerol were selected as potential ligands. Tests using commercial OA and glycerol showed that NlGr23a-Sf9 cells responded dramatically to OA stimulation (Figure 1E).

**Figure 1.**
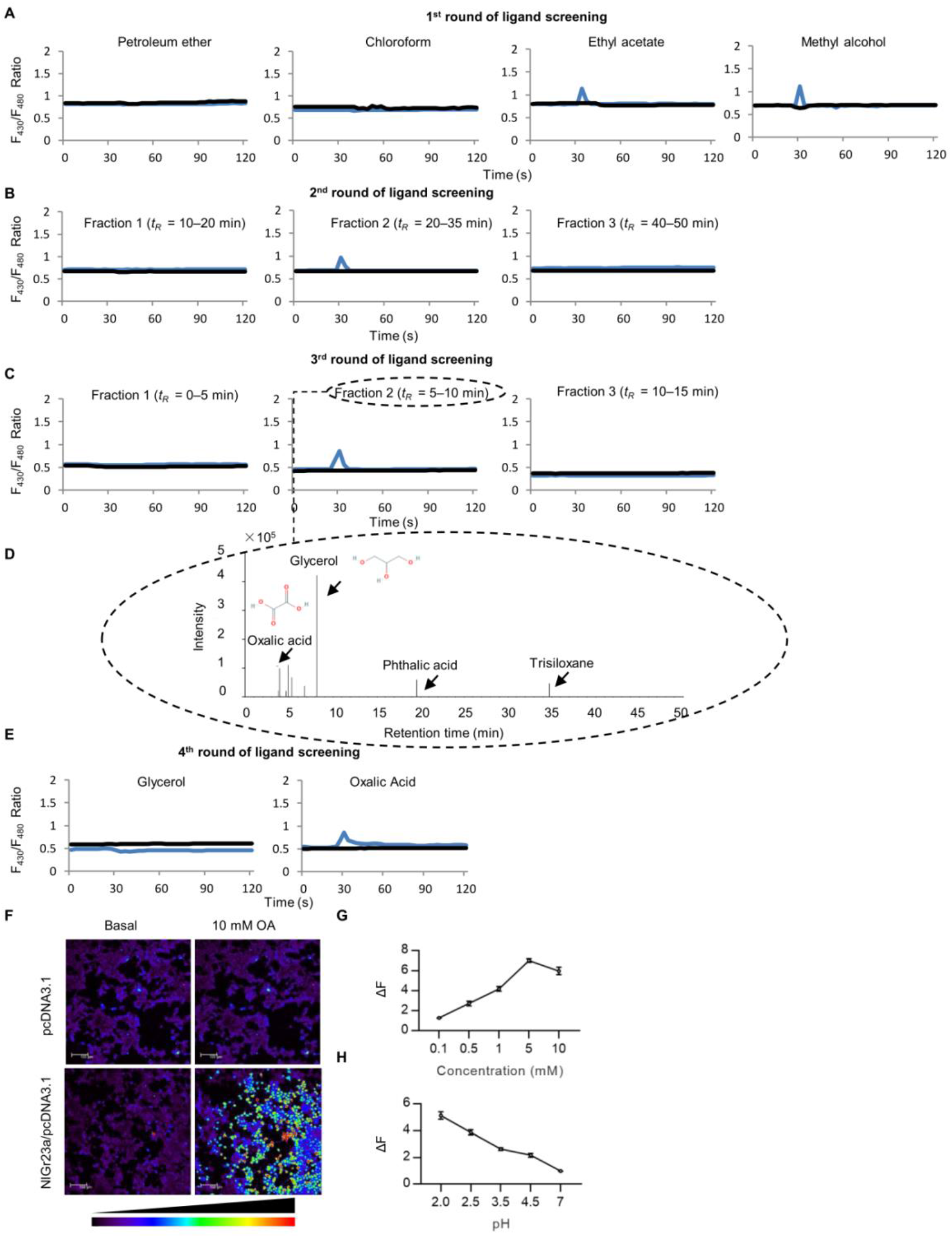
Oxalic acid is a ligand of NlGr23a. (A) Round 1 of ligand determination. The responses of Sf9 cells transfected with plasmid construct pIZ-NlGr23a-V5-His (blue lines) or pIZ-V5-His (black lines) vector to different fractions and tastants were analysed by calcium imaging. The changes in intracellular Ca^2+^ indicated by the ratio of F_340_/F_380_ in Fura-2 AM loaded cells in response to fractions of the crude extract obtained with petroleum ether (PE), chloroform (CHCl_3_), ethyl acetate (EAC) and methyl alcohol (MeOH) were measured using microscopy. Test solutions are shown above each figure. Each curve represented one cell responding to a specific tastant. (B) Round 2 of ligand determination. Representative traces evoked by three semi-preparative high-performance liquid chromatographic (semi-prep-HPLC) fractions separated from the EAC fraction. See Appendix 1**—**figure 2A for HPLC chromatogram of the EAC fraction. (C) Round 3 of ligand determination. Representative traces evoked by three semi-prep-HPLC fractions separated from the bioassay-positive fraction from round 2 of ligand screening. See Appendix 1**—**figure 2B for HPLC chromatogram of the bioassay-positive subfraction. (D) GC-MS chromatogram of bioassay-positive fraction from round 3 of ligand screening. Arrowheads indicate substances identified in the bioassay-positive fraction. See Appendix 1**—**figure 2C for GC-MS chromatogram of the control solution and Appendix 1**—**table 1 for the retention time and response intensity of each compound. Chemical structure of oxalic acid (OA, PubChem CID: 971) and glycerol (PubChem CID: 753) were obtained from *PubChem* (https://pubchem.ncbi.nlm.nih.gov). (E) Round 4 of ligand determination. Representative traces evoked by 5 mM glycerol and 5 mM OA. (F) Cytoplasmic Ca^2+^ imaging of the HEK293T cells expressing NlGr23a. Upper: the pcDNA3.1 transfected HEK293T cells. Lower: the NlGr23a/pcDNA3.1 transfected HEK293T cells. Left: Fluorescence image before 10 mM OA perfusion. Right: Fluorescence image after 10 mM OA perfusion. White scale bar indicates 100 μm. The color scale indicates the cytoplasmic fluorescence ratio. (G) Ca^2+^response of the NlGr23a-HEK293T cells (*n* = 50) stimulated with various concentrations of OA solutions. ΔF represented the values of F_340_/F_380_ ratio, which were set to 1 before the addition of the OA solution. Bars represent means ± SEM. (H) Ca^2+^response of the NlGr23a-HEK293T cells (*n* = 50, except for pH = 4.5, where n = 27) stimulated with 10 mM OA at different pH. ΔF represented the values of F_340_/F_380_ ratio, which were set to 1 before the addition of the compound. pH was adjusted using NaOH. Bars represent means ± SEM.

**Figure 2.**
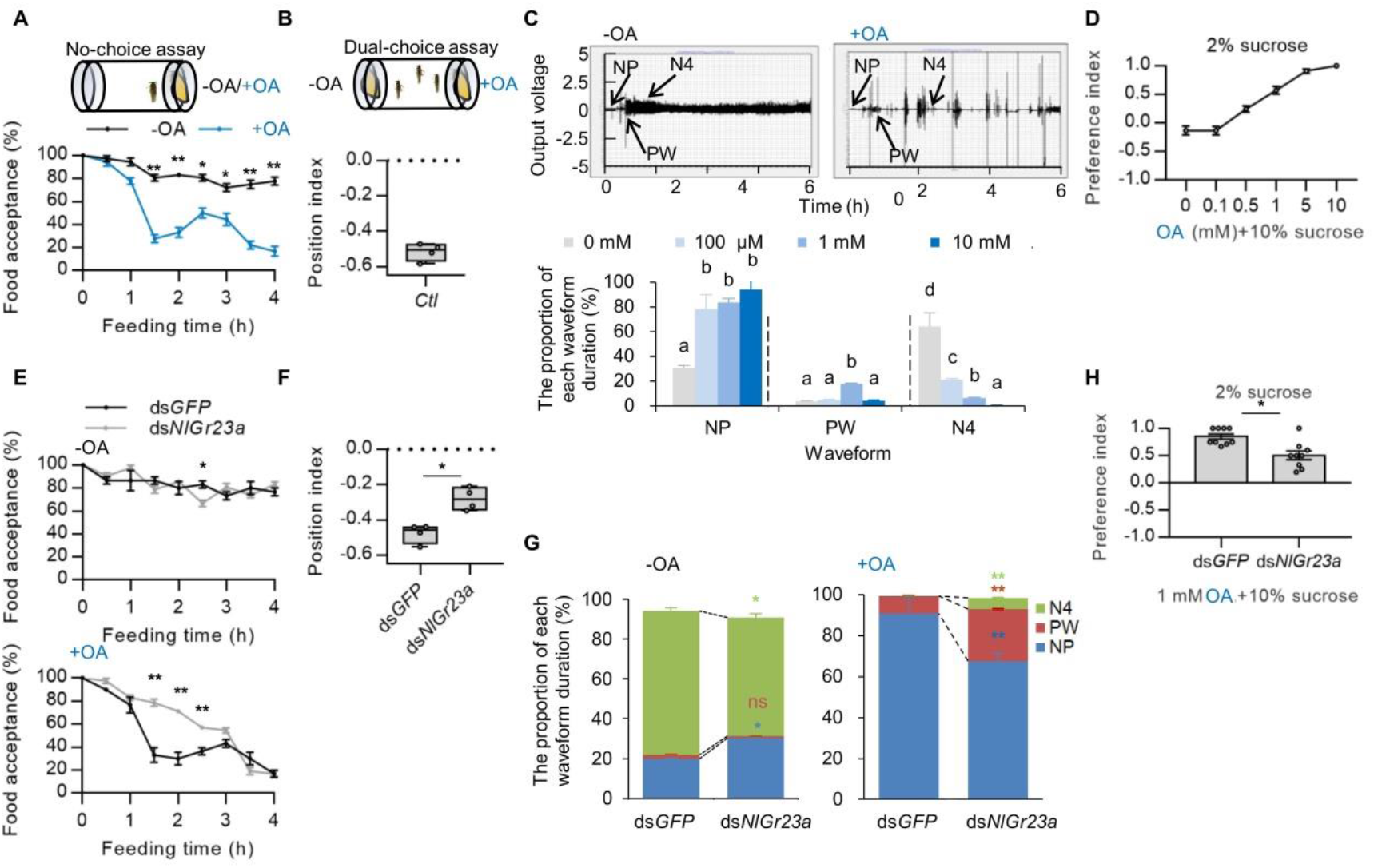
Effects of *NlGr23a* on the antifeedant activity of oxalic acid. (A) Top: Schematic representation of the experimental apparatus used for no-choice test. Bottom: The proportion of BPHs accepting artificial diets without (-OA, in black) or with 100 μM OA (+OA, in blue) as food in no-choice test. The data represent means ± SEM. *t*-test; \**P* < 0.05; \*\**P* < 0.01 (*n* = 3). (B) Top: Schematic representation of the experimental apparatus used for dual-choice test. Bottom: Collective data (*n* = 4) of the position index (PI) of the laboratory insects (*Ctl*) in response to food containing 100 μM OA in dual-choice assays. (C) Top: Overall typical view of electrical penetration graph (EPG) waveforms generated by the feeding behaviours of BPH on liquid diet sacs (LDS) without (top) or with (bottom) 100 μM OA. There are three typical waveforms: non-penetration (NP), pathway (PW), and ingestion (N4). Bottom: The duration of each EPG waveform as a proportion of observation time produced by BPHs on LDS containing artificial diets with different concentrations of OA. Bars represent means + SEM. Different lowercase letters above columns represent significant differences at *P* < 0.05 from Duncan’s multiple range test (*n* = 5). (D) Dose-responsive food choice assay for OA. BPHs were given the choice between 2% sucrose solution and 10% sucrose with different concentrations of OA. The data represent means ± SEM (n = 8). (E) Top: The proportion of insects accepting artificial food in absent of OA (-OA) among ds*GFP*-treated (black line) or ds*NlGr23a-* treated (grey line) BPHs in no-choice tests. Bottom: The proportion of insects accepting artificial food with 100 μM OA (+OA) among ds*GFP*-treated or ds*NlGr23a-* treated BPHs in no-choice tests. The data represent means ± SEM. *t*-test; \**P* < 0.05; \*\**P* < 0.01 (*n* = 3). (F) PI values of ds*NlGr23a*-treated and ds*GFP*-treated BPHs in response to +OA diets in dual-choice assays. *t*-test; \**P* < 0.05 (*n* = 4). (G) The duration of each EPG waveform as a proportion of observation time produced by ds*GFP*-treated and ds*NlGr23a*-treated BPHs on LDS with or without 10 mM OA. Stacked bar graphs represent means + SEM. *t*-test; ns, *P* > 0.05; \**P* < 0.05; \*\**P* < 0.01 (*n* = 5). (H) Food choice assay for ds*NlGr23a*-treated and ds*GFP*-treated BPHs. 1 mM OA was mixed with 10% sucrose. Bars represent means ± SEM. *t*-test; \**P* < 0.05 (n ≥ 7).

In addition, expressing NIGr23a in the human embryonic kidney 293T (HEK293T) cells also allowed these cells to respond to OA. This suggests that NIGr23a showed full functionality in Sf9 cells without an insect-specific co-receptor (Figure 1F). The responses elicited by the HEK293T cells increased with higher acid concentrations and were inversely correlated with pH of OA solution (Figs 1G and H). Microscale thermophoresis assay, which has been used to measure binding affinities, supported the results that NlGr23a interacts with OA. As seen in Figure 1—figure supplement 1A, when NlGr23a-GFP was mixed with different concentrations of glycerol, there was no fluorescence migration change after laser irradiation, suggested that the binding state of NlGr23a did not change by glycerol, whereas when NlGr23a-GFP was mixed with different concentrations of OA, the fluorescence migration via laser irradiation gradually decreased as the concentration of OA increases. Thus, OA was shown to be a ligand of NlGr23a. As NlGr23a interacted with bioactive molecules in the MeOH fraction, NlGr23a may also have non-OA ligands in rice (Figure 1A). To characterize the response profiles of NlGr23a, we tested responsiveness of the NlGr23a-expressing HEK293T cells to ten additional phytochemicals. Stimulation with only benzoic acid induced a Ca^2+^ increase, whereas stimulation with other organic acids, which have shown inhibitory effects against BPHs, elicited no response (Yoshihara *et al.*, 1980) (Figure 1—figure supplement 1B). No response was also observed in sucrose, various bitter compounds and HCl (Figure 1—figure supplement 1B). Hence, NlGr23a may mediate carboxylic acids sensing in a structure dependent manner.

### *NlGr23a* is essential to the antifeedant activity of oxalic acid

BPH adults offered artificial diets either containing (+OA) or missing OA (-OA) with no alternative (“no-choice” tests) showed significantly lower acceptance of +OA compared to -OA after 1.5 h exposure (Figure 2A). When offered a choice (“dual-choice”), +OA food was avoided in favour of -OA food, based on insect positioning relative to the food sources (position index, PI = −0.52, Figure 2B). The electrical penetration graph (EPG) technique, which can discriminate between feeding behaviours by producing different waveforms, was used to investigate the effect of OA on BPH feeding behaviour. Three such waveforms occurred sequentially during the process of BPH feeding on liquid diet sacs (LDS): NP (non-penetration), PW (pathway wave), and N4 (artificial diet ingestion) (Figure 2C, top). At 100 μM OA in the diet (+OA), the pre-feeding phases were longer than those of the control (-OA) and the N4 phase was very significantly shorter (Figure 2C, bottom). OA, even at a low concentration, was thus capable of interfering food sucking behaviour. The proportion of time spent feeding fell further as OA concentrations increased, down to only 0.56% at an OA concentration of 10 mM (Figure 2C, bottom).

Further, we added two different food dyes into sucrose solutions with or without OA to measure direct feeding. BPHs were given a choice between 2% (w/v) sucrose and 10% sucrose plus different concentrations of OA. Sucrose is a potent sucking stimulant for BPH (Sogawa, 1982). In the absence of OA, the insects showed a preference for the 5-fold higher concentration of sucrose (Figure 2D). However, when the higher concentration of sucrose was contaminated with OA, BPHs avoided the OA-laced food in a dose dependent manner (Figure 2D). A similar experiment was conducted using an equal concentration of sucrose (10%), BPHs showed similar preference for 10% sucrose solutions added with different food dyes without OA and exhibited repulsion to the food when it was mixed with OA (Figure 2—figure supplement 1A). In addition, BPHs consumed significantly less food containing much less concentration of OA (100 μM) compared to the control (Figure 2—figure supplement 1B). These results indicated that the antifeedant activity of OA was dose dependent. To exclude the possibility of olfaction mediated OA sensing, we surgically removed the primary olfactory organs of BPH, the second and third antennal segments (Aljunid and Anderson, 1983). Antennaectomized insects were normal for OA avoidance (Figure 2—figure supplement 1C). Thus, olfactory system is not required for OA avoidance.

As OA was a ligand of the NlGr23a receptor, we hypothesised that the antifeedant activity of OA is dependent on this receptor. The *NlGr23a* gene was therefore silenced using RNA interference (RNAi) *in vivo*. *NlGr23a* expression in the whole insect was decreased significantly 24 to 72 h post-injection (Figure 2—figure supplement 2A). Additionally, *NlGr23a* mRNA levels in primary gustatory organs at 48 h post injection substantially decreased compared to the negative controls (Figure 2—figure supplement 2B). As the tissue amount of tarsus was insufficient for RNA extraction, we extracted RNA from legs to detect the gene silencing efficiency. As expected, the antifeedant activity of OA was decreased in *NlGr23a*-inactive insects compared to the control insects injected with green fluorescent protein (*GFP*) double-stranded RNA (dsRNA) in no-choice tests. In BPHs fed -OA artificial diets, the level of food acceptance was nearly the same for both treated (ds*NlGr23a*) and control (ds*GFP*) groups (Figure 2E, top). In insects fed +OA diets, those with silenced *NlGr23a* spent more time on the OA-containing food than controls, significantly so between 1.5 h and 2.5 h after exposure (Figure 2E, bottom). In the dual-choice assays, a decrease in OA’s antifeedant effectiveness was also observed. BPHs injected with ds*NlGr23a* were less sensitive to OA (PI=−0.28) than controls (PI=−0.48, Figure 2F). When BPHs fed on diets without OA, the NP duration was higher and the N4 duration was commensurately lower in *NlGr23a*-silenced insects (Figure 2G, left), which indicated that RNAi against *NlGr23a* influenced their feeding behaviour. At 10 mM OA, *NlGr23a*-silenced BPHs exhibited significantly more N4 and PW phases and fewer NP phases than controls (Figure 2G, right). Similar results were observed at 2 mM or 5 mM OA (Figure 2—figure supplement 2C). Additionally, knockdown of *NlGr23a* dramatically reduced OA feeding avoidance (Figure 2H). Taken together, these results indicated that *NlGr23a*-silenced BPHs were less sensitive to OA.

### *NlGr23a* is required for perception of oxalic acid in rice

To further verify above results in rice plants, we used the immersion method to increase the OA content of the rice variety TN1, resulting in 40% higher content in the OA-treated rice plants (TN1^+OA^) than in the untreated controls (Figure 3A). Insects fed TN1^+OA^ showed a positional avoidance response (PI =−0.24) and made twice as many probing marks (caused by insects testing food for palatability and withdrawing) as those fed TN1, indicating that BPHs spent more time searching for food on TN1^+OA^ (Figs 3B and 3D). There were also three main types of EPG waveforms occurred during the process of BPH feeding on a rice stem: NP (non-penetration), PW (pathway wave) and N4 (phloem ingestion). Compared to the control group reared on TN1, BPHs fed TN1^+OA^ exhibited longer PW and shorter N4 durations, and thus less phloem ingestion (Figure 3F). Our analysis therefore indicated that BPH feeding was inhibited as the content of OA in rice plants increased.

**Figure 3.**
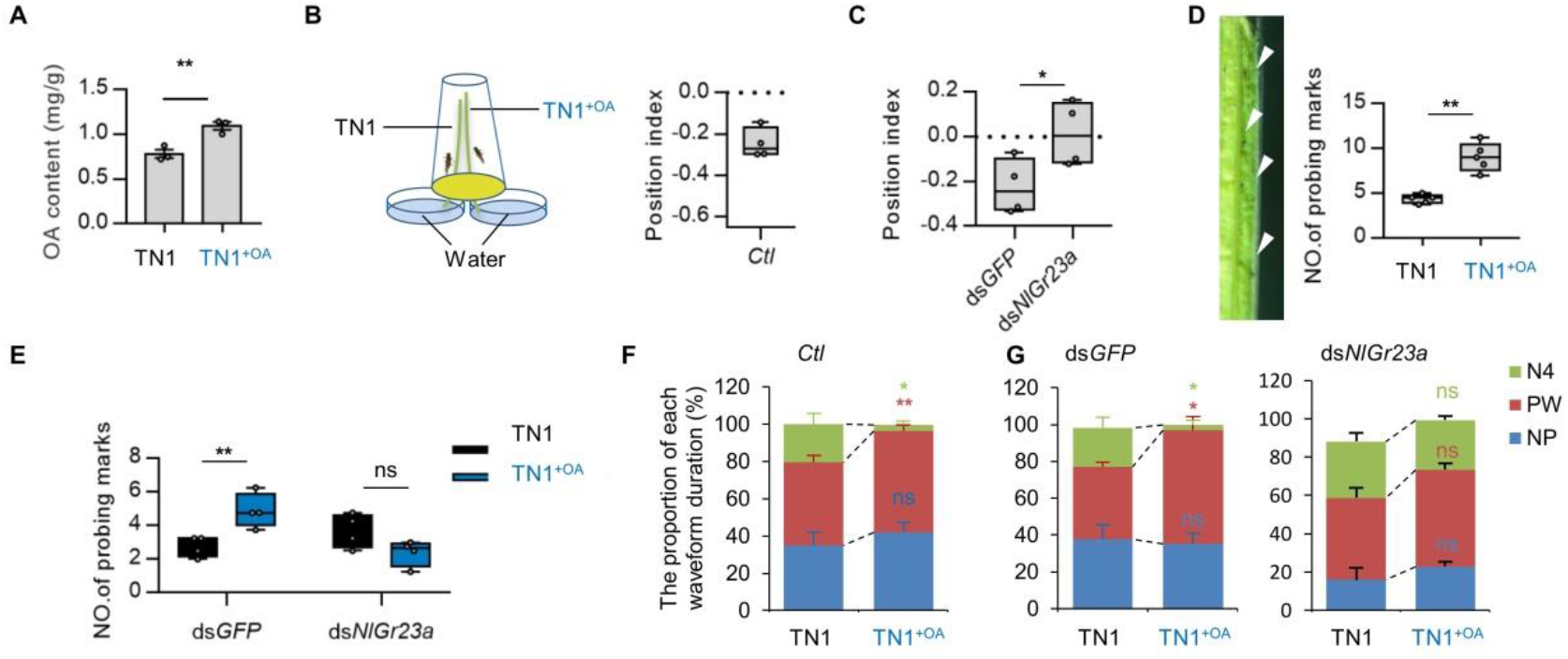
Increased OA content in rice plant inhibited feeding behaviour of BPH. (A) OA content before and after immersion treatment in TN1. The TN1^+OA^ represented TN1 treated with 0.2% OA solution. Bars represent means ± SEM. *t*-test; \*\**P* < 0.01 (*n* = 4). (B) Left: Schematic representation of the experimental apparatus used for dual-choice test with rice plants. Right: Collective data (*n* = 4) of the position index (PI) of *Ctl* in response to TN1^+OA^. (C) PI values for *NlGr23a* dsRNA-treated and *GFP* dsRNA-treated BPHs in response to TN1^+OA^ in dual choice assays. *t*-test; \**P* < 0.05 (*n* = 4). (D) Left: Probing marks left on the surface of a rice stem. Right: Collective data (*n* = 5) of the number of probing marks on the surface of TN1^+OA^ or TN1 after BPH infestation. *t*-test; \*\**P* < 0.01. (E) Collective data (*n* = 4) of the number of probing marks on the surface of TN1^+OA^ or TN1 made by BPHs treated with *NlGr23a* dsRNA or *GFP* dsRNA. *t*-test; ns, *P* > 0.05; \*\**P* < 0.01. (F) The duration of each EPG waveform as a proportion of observation time produced by BPHs on TN1 or TN1^+OA^. Stacked bar graphs represent means + SEM. *t*-test; ns, *P* > 0.05; \**P* < 0.05; \*\**P* < 0.01 (*n* = 5). (G) The duration of each waveform as a proportion of observation time produced by *NlGr23a*-silenced and *GFP* dsRNA-treated BPHs on TN1 or TN1^+OA^. Bars represent means + SEM. *t*-test; ns, *P* > 0.05; \**P* <0.05.

The positional avoidance response disappeared in ds*NlGr23a*-treated BPHs (PI = 0.01) but not in ds*GFP*-treated insects (PI = −0.22) (Figure 3C). In addition, ds*GFP*-treated BPHs produced 77% more probing marks on TN1^+OA^ rice plants than on TN1, but this difference also disappeared after injection of *NlGr23a* dsRNA (Figure 3E). EPG results further supported this phenomenon. The proportion of time in the N4 waveform was significantly lower and PW waveform was significantly higher when BPHs fed on TN1^+OA^ in ds*GFP*-treated group, but this significant difference no longer existed after *NlGr23a* was silenced (Figure 3G). These results further confirmed that *NlGr23a* mediated OA perception in rice.

As we observed, the PI values in dual choice tests in rice tended to be higher than those from tests using the artificial diet containing OA (Figs 2B and 3B). There are three explanations. Firstly, in the case of the artificial diets, the diets either included or completely excluded OA, while in the case of rice plants, the OA content in TN1^+OA^ was only 39% higher than in TN1 (Figure 3A). Secondly, BPHs tend to move up and down along the same plant instead of quickly moving to a different plant. Thirdly, rice plants contain many metabolites including both deterrents and stimulants to feeding (Yoshihara *et al.*, 1980; Zhang *et al.*, 1999).

### *NlGr23a* is also indispensable for sensing OA via BPH tarsi

Given that *NlGR23a* was highly expressed in the tarsus (Figure 4A), we developed a novel experiment referred to a previous method (Lee *et al.*, 2010), tarsal avoidance response (TAR) assay, to explore whether *NlGr23a* functioned in OA detection via the tarsi (see Material and Method for the detail of TAR assay). We first determined the behavioral significance of OA to the tarsal behavior by presenting sucrose or sucrose plus OA as a stimulus to the legs. The tarsi preferably adhere to Kimwipe dipped in sucrose (Figure 4B and see also Figure 4—video supplement 1). However, the tarsi elicited TAR after adding OA into sucrose solution (Figure 4B and see also Figure 4—video supplement 2). A small increase was observed in TAR in the second exposure of sucrose alone, likely due to the residual OA effects. Moreover, we observed that OA solution triggered a TAR in a dose dependent manner (Figure 4C). This experiment clearly indicates that female BPH is equipped with a sensitive detection system for OA in the tarsi. As expected, ds*NlGr23a*-injected BPH exhibited a significant reduction in TAR compare to the control (Figure 4D), which supported the idea that *NlGr23a* is indispensable for sensing OA via BPH tarsi.

**Figure 4.**
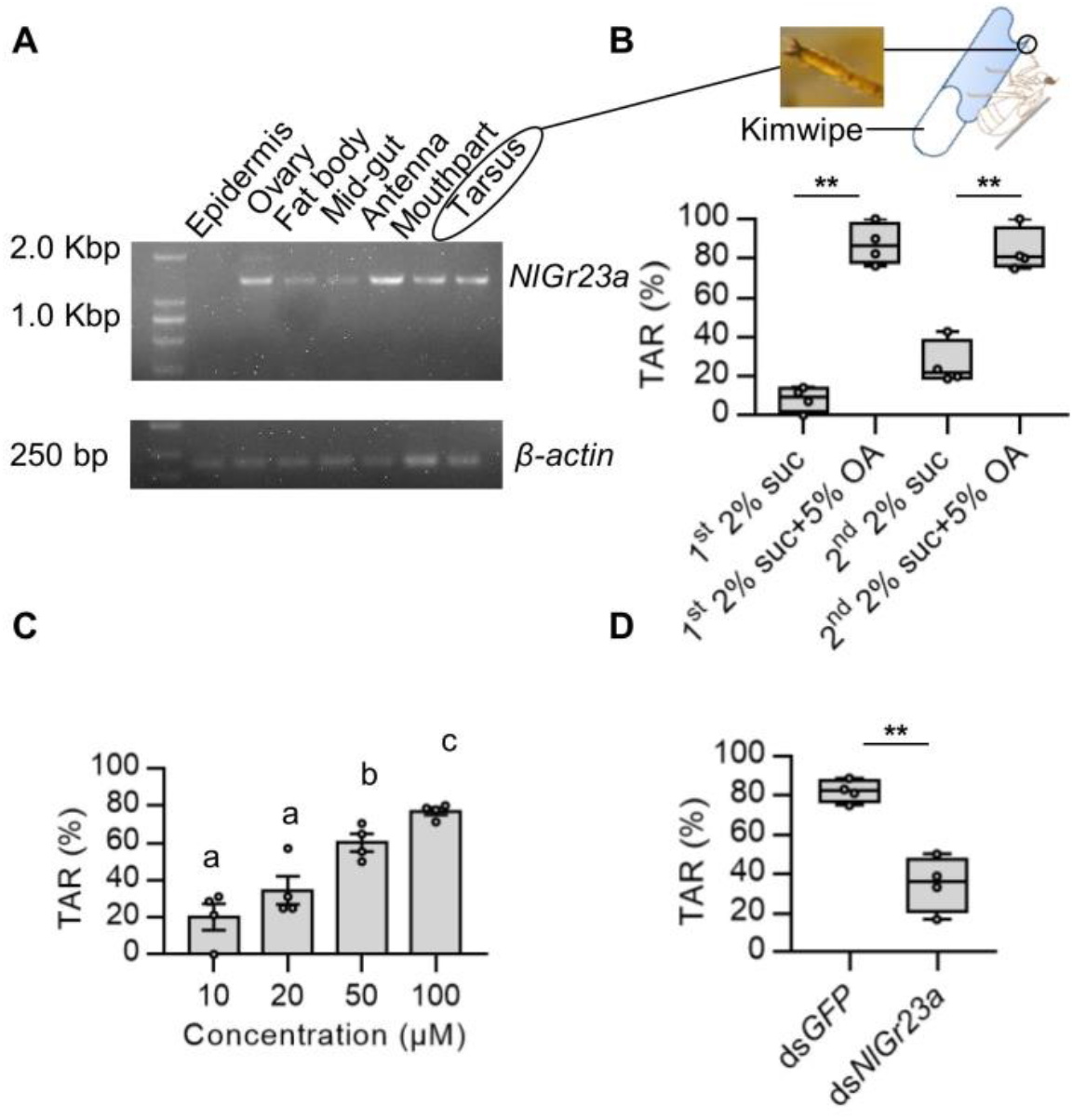
*NlGr23a* is indispensable for sensing OA via BPH tarsi. (A) RT–PCR of *NlGr23a* on RNA isolated from various organs and tissues. All samples were isolated from one-day-old brachypterous female adults. (B) Top: Schematic illustration of tarsal avoidance response (TAR) assay for BPHs. Bottom: Collective data (*n* = 4) of TAR of BPHs in response to different stimulants. Insects were initially given 2% sucrose, then 2% sucrose in combination with 5% OA. The trial was conducted two rounds. *t*-test; \*\**P* < 0.01. (C) TAR in BPH upon legs stimulation by 10, 20, 50, 100 μM solutions of OA. Bars represent means ± SEM. Different lowercase letters above columns represent significant differences at *P* < 0.05 from Duncan’s multiple range test (*n* = 4). (D) TAR values for *NlGr23a* dsRNA-treated and *GFP* dsRNA-treated BPHs in response to 100 μM OA. *t*-test; \*\**P* < 0.01 (*n* = 4). **Figure 4—video supplement 1.Tarsal avoidance response (TAR) to sucrose.** A Kimwipe containing 2% sucrose was applied to the distal part of legs of BPH. BPH showed acceptance towards the wet wick and its tarsi preferably adhere to Kimwipe. **Figure 4—video supplement 2. Tarsal avoidance response (TAR) to oxalic acid.** A Kimwipe containing 2% sucrose plus 5% oxalic acid was applied to the distal part of legs of BPH (the same as the one in Figure 4—video supplement 1) at 5 min interval after 2% sucrose treatment. BPH refused to adhere to the wick and used leg to push away Kimwipe. The hindlegs of BPH did not contact Kimwipe.

## Discussion

Although insect bitter Grs are assumed to detect plant secondary compounds or bitter tastants (Robertson and Wanner, 2006), it is still very difficult to identify the ligands of bitter Grs, especially new examples. The pioneering identifications typically used electrophysiological and behavioural genetic analyses in Gr mutants of the model insect *D. melanogaster*. Additionally, nearly all of the ligands tested to date have been directly purchased compounds such as caffeine, strychnine, umbelliferone, saponin, chloroquine, and L-canavanine (Moon *et al.*, 2006; Poudel *et al.*, 2015; Shim *et al.*, 2015; Poudel *et al.*, 2017; Sang *et al.*, 2019). In other species, especially in agricultural insect pests, Gr ligands are often assumed to match those of the orthologous fruit fly receptors. As non-sugar *Gr* genes are subject to rapid adaptation driven by the vastly different ecological niches occupied by insect species and shared sequence identity among species is low, it is difficult to identify matching receptors. Alternatively, candidate ligands have been chosen from chemical classes that are known to elicit neurophysiological or behavioural responses. For example, a collection of known oviposition stimulants of *Papilionidae xuthus* were selected to test for interaction with PxutGr1, and eventually synephrine was identified as the specific ligand (Ozaki *et al.*, 2011).

In order to expand potential ligand sources for non-sugar Grs, plant extracts have been tested in few cases. Extracts of whole citrus caused an increase in Ca^2+^-dependent luminescence in Sf9 cells after the introduction of *PxutGr1* (Ozaki *et al.*, 2011), and similar results were observed for crude extracts of cotton leaves and 3 *Grs* in *Helicoverpa armigera* (*HarmGr35*, *HarmGr50*, and *HarmGr195*) (Xu *et al.*, 2016). However, no specific ligand has so far been identified based on separation of these crude extracts. In this study, we started from rice crude extracts and eventually identified OA as a ligand of NlGr23a. To our knowledge, OA is the first identified ligand starting from plant crude extracts, and the first strong acidic ligand for insect Gr proteins.

Grs in insects are known to form heterodimers with co-receptors to recognize the tastant (Dahanukar *et al*., 2007; Lee *et al*., 2009). Nevertheless, there are several instances of the expression of a single insect Gr in a heterologous expression system sufficient to confer tastant responses, including Gr5a, Gr43a and Gr64e from *Drosophila*, BmGr8 and BmGr9 from the silkworm *Bombyx mori*, PxutGr1 from the swallowtail butterfly *Papilio xuthus*, HarmGR35, HarmGR50 and HarmGR195 from the polyphagous pest *Helicoverpa armigera*, AgGr25 from the malaria mosquito *Anopheles gambiae* (Chyb *et al*., 2003; Freeman *et al*., 2014; Sato *et al*, 2011; Zhang *et al*., 2011; Ozaki *et al*., 2011; Xu *et al*., 2016), and PrapGr28 from the cabbage butterfly *Pieris rapae* (Yang *et al*., 2021). In this study, the HEK293T cells expressing NlGr23a caused responses to OA (Figure 1F). But we do not exclude a possibility that NlGr23a requires co-receptors for its fully functionality *in vivo*. Based on the analysis tuned to a 7TMPs characteristic of insect Grs and a homology search with known broadly required gustatory co-receptors, NlGr2, NlGr11, NlGr13, NlGr15, NlGr17, NlGr18, NlGr21 and NlGr26 were selected for food preference assay with 1 mM OA (Kang *et al*., 2018). Gene silencing of the target genes was achieved by dsRNA injection (See Appendix 3—figure 1A for the silencing efficiency of target genes and Appendix 2—table 1 for primers used for dsRNA synthesis and RNAi efficiency detection). Those BPHs treated with ds*NlGr2* and ds*NlGr13* were less sensitive to OA than controls (Appendix 3—figure 1B). These results suggest that NlGr2 and NlGr13 might act as co-receptors of NlGr23a for OA recognition or through NlGr23a-independent mechanisms.

OA is, however, commonly present in rice plants. The species of only 11 out of 93 orders of higher plants do not store OA (Zindler-Frank, 1976). The fact that a secondary compound is deterrent to an insect does not imply that the insect does not eat plants containing that compound. Whether or not it does so depends on its sensitivity to the compound, the background of other deterrent and phagostimulatory information in which it is perceived, and the degree of food deprivation incurred by the insect (Chapman, 2003). In the EPG experiments of this study, OA concentrations in artificial diets and in rice plants significantly inhibiting feeding phases were 1 mM and about 10 mM (the OA content of the background material TN1 was measured as above 5 mM), respectively (Fig.2C, Fig. 3A and F). The difference in active OA concentration may be due to the different background of other chemicals. Moreover, OA concentrations in rice plants are dynamic, maybe varied in tissues, and affected by rhythm, rice variety and developmental stages, as well as abiotic factors (Wu *et al.*, 2016). OA isolated from leaf sheath extracts of rice seems to mediate the phloem-feeding habit as a general inhibitor which commonly occurs in plant tissues outside the phloem (Yoshihara *et al.*, 1980). We also measured OA contents in three rice varieties and indeed found there is a significant difference in OA content between the susceptible TN1 rice and the resistant rice varieties IR36 and IR56 (Appendix 4—figure 1). Thus, we speculate that a population of sensory neurons tuned to OA is probably involved in detection of the dynamic changes of soluble OA to provide clues for the suitable habitats or feeding times.

Insects discriminate a wide range of tastants, including sugars, bitter compounds, NaCl, and sour substances. In contrast to sweet and bitter-tasting chemicals, acids elicit varied behavioural responses, depending on their structure and concentration (Rimal *et al.*, 2019). In *Drosophila*, both Ir25a and Gr64a-f expressed in sweet taste neurons mediated appetitive responses to lactic acid (Stanley *et al*., 2021). Knock-down of *Ir25a*, *Ir76b* and *Ir56d* abolished fatty acid recognition (Ahn *et al*., 2017). By screening individual Gr64 cluster gene mutant flies, Gr64e was also identified as a receptor mediating fatty acid sensing (Kim *et al.*, 2018). Ir7a expressed in bitter GRNs is required for rejecting foods laced with high concentrations of acetic acid (Rimal *et al*., 2019). So far, most chemoreceptors repsond to sour tastes are ionotropic receptors, such as Ir25a and Ir76b (Chen and Dahanukar, 2020). In this study, we investigated an important plant-insect interaction and found that NlGr23a in BPH senses OA in rice.

It was reported that behavioural repulsion from acidic stimuli is mediated by two different classes of GRN in the fruit fly: those sensing sugar and bitters (Charlu *et al.*, 2013; Devineni *et al.*, 2019). This implies that Grs sensing sour compounds may influence insect behaviour through multiple pathways. After *NlGr23a* was silenced, BPHs with *NlGr23a* dsRNA were used for transcriptome and KEGG pathway analysis of the differentially expressed genes. These analyses showed that NlGr23a may function through diverse signalling pathways, including mTOR, AMPK, insulin, and PI3K-AKT (see Appendix 5—figure 1). The possible molecular mechanisms of NlGr23a in modulating feeding behaviour will be further investigated.

## Materials and Methods

### Insect and rice culture

A BPH laboratory strain (*Ctl*) was obtained from Guangdong Academy of Agricultural Sciences (GDAAS, Guangdong, China), and this strain was reared in a continuous laboratory culture on susceptible rice seedlings (variety Huang Hua Zhan). All BPHs were maintained at 26 ± 2 °C with 80 ± 10% humidity and a light-dark cycle of L16:D8 h. One-day-old brachypterous female adults were collected for the following bioassays. Plants aged 30–40 days of the susceptible rice variety TN1 were used for the relevant bioassays. Allocation of insects was randomly done to minimize the effects of subjective bias.

### Chemical sources

OA, glycerine, adonitol, methoxylamine hydrochloride, *N*-methyl-*N*-(trimethylsilyl) trifluoroacetamide, tetrabutylammonium bisulphate (TBA), KH_2_PO_4_ and strychnine and caffeine (purity, ≥99%) were purchased from Sigma-Aldrich Company (St. Louis, MO, USA). HPLC grade reagents including petroleum ether (PE), chloroform (CH_3_Cl), ethyl acetate (EAC), methyl alcohol (MeOH), acetonitrile (ACN), isopropyl alcohol and pyridine, and analytical grade reagents including sucrose, trans-aconitic acid, benzoic acid, salicylic acid, mandelic acid, quinine and ethyl alcohol were all obtained from Aladdin Reagent (Shanghai, China). Maleic acid was purchased from MedChemExpress (Shanghai, China). Hydrochloric acid (HCl) was provided by the Guangzhou Chemical Reagent (Guangzhou, China).

### Ligand determination for NlGr23a

#### *NlGr23a* cDNA cloning and stable cell line construction

Total RNA from BPHs was prepared as previously described (Kang *et al.*, 2018). A PrimeScript™ RT reagent kit with gDNA Eraser (Takara, Kyoto, Japan) was used with 1 μg RNA for first -strand complementary DNA (cDNA) synthesis. A fragment of full-length *NlGr23a* cDNA was amplified using 2×SuperStar PCR Mix with Loading Dye (Genstar, Beijing, China) following the manufacturer’s instructions. The cloning primers are listed in Appendix 2—table 1.

The *NlGr23a* cDNA was inserted into the Hind III and EcoR I sites of pIZ/V5-His vectors (Invitrogen, Carlsbad, CA, USA) using T4 DNA ligase (NEB, Beijing, China). The NlGr23a-Sf9 stable cell lines were obtained by using a previously described method (Chen *et al.*, 2021). Briefly, Sf9 cells were cultured in Grace’s Insect Medium (Gibco, Grand Island, NY, USA) supplemented with 10% foetal bovine serum at 27 °C. Cells were plated into 6 -well plates and left to settle for 20 min before being transfected with either 2 μg of the recombinant plasmid pIZ-NlGr23a-V5-His or pIZ-V5-His vector (negative control), and 6 μL Fugene HD transfection reagent (Promega, Madison, WI, USA) in 100 μL per well of Grace’s Insect Medium. Forty-eight hours after transfection, cells were cultivated under Zeocin selection. The final concentration of Zeocin to maintain cells was 100 μg/mL. Following antibiotic selection for at least 10 passages, stable cell lines were verified by immunoblotting and RT-PCR. For western blotting analysis, cells were lysed by total membrane proteins extraction buffer A (Invent Biotechnologies, Inc., Eden Prairie, USA). The supernatant of the mixture was further centrifuged. The sediment contains the total membrane proteins while the supernatant contains cytosol proteins. Samples were then immunoblotted with anti-V5 antibody. For RT-qPCR, we extracted total RNA of cells after antibiotic selection for cDNA template synthesis. PCR was performed under the following conditions: 94°C for 2 min, 35 cycles of 94°C for 30 s, 55 °C for 30 s, 72°C for 90 s, and 72°C for 5 min. PCR amplification products were run on a 1.2% agarose gel.

The HEK293T cells were cultured in Dulbecco’s modified Eagle medium (Gibco, California, USA) supplemented with 10% fetal bovine serum in an atmosphere of 5% CO_2_ and 95% relative humidity at 37°C. To confirm the sufficiency of NIGr23a for OA-mediated response, the HEK293T cells were transiently transfected with NlGr23a/pcDNA3.1-FLAG. NlGr23a/pcDNA3.1-FLAG was constructed by inserting a C-terminally FLAG-tagged human codon-optimized *NlGr23a* ORF into the pcDNA3.1 vector using BamHI and EcoRI sites. The nucleotide sequences of the codon-optimized genes were listed in Appendix 6—figure 1. Each 0.5 μg plasmid was transfected into the HEK293T cells using Lipofectamine 3000 (Invitrogen, Carlsbad, CA, USA), according to the manufacturer’s instruction.

#### Isolation and identification of active compounds in rice

A bioassay-guided approach was adopted to isolate and identify active compounds interacting with NlGr23a from the crude extracts of rice plants. Firstly, we ground stems and leaves of TN1 (5 g), and performed serial solvent extractions on macerated subsamples with 25 mL of isopropanol (IPA) at 75 °C for 15 min, 25 mL CHCl _3_–IPA (1:2, v/v; 5 h; room temperature), and 25 mL MeOH–CHCl_3_ (2:1, v/v; overnight; room temperature). Each extract was centrifuged at 2,000 rpm for 15 min to collect the supernatant. The pooled supernatants were then concentrated to dryness in vacuum at 25 °C and re -dissolved in 12 mL CHCl_3_ to obtain the crude rice extract.

Secondly, 6 mL crude extract was subjected to a column chromatography with silica gel, eluting with solvents in the order of increasing polarity (PE - CHCl_3_ - EAC - MeOH) to yield four fractions. We carried out the first round of ligand screening using these four initial fractions. Thirdly, semi-preparative HPLC separation of the bioassay-active fraction I was performed on an Agilent 1260 HPLC system (Agilent Technologies, Santa Clara, CA, USA) equipped with an Agilent ZRBAX SB-C18 column (4.6 mm × 150 mm, 3.5 μm) by gradient elution with ACN (A) and water (B). The gradient program was as follows: 3 min, 5% B; 22 min, 5–20% B; 15 min, 20-40% B; 15 min, 40–50% B; 8 min, 50–95% B; 12 min, 95-5% B, all at a flow-rate of 1 mL/min. The eluate from the column was continuously monitored at 205 and 290 nm after sample loading (5 μL), and the fraction collector was programmed to collect fractions during periods in which most substances were detected (see Appendix 1—figure 2A). The second round of screening for the ligand was then carried out with these subfractions. Fourthly, the active semi-prep-HPLC fraction (active fraction II) was subjected to further fractionation with 100% ACN at a flow-rate of 1 mL/min for the third round of screening. Last, the bioassay-positive fraction from this latest round (active fraction III) was derivatized and elucidated using GC-MS. The GC-MS detection for the derived samples was performed on an Agilent 7890A gas chromatograph equipped with an Agilent 5975C VL MSD detector (Agilent Technologies, Santa Clara, CA, USA) as previously described (Kang *et al.*, 2019).

#### Calcium imaging assay

Calcium imaging using fura-2/AM calcium indicator dye was performed as previously described with some modification (Nelson *et al.*, 2001; Chen *et al.*, 2021). Cells were seeded in a glass bottom cell culture dish (φ20 mm, Nest, Jiangsu, China) at 70% confluency and grown overnight. After being washed in Hanks’ balanced salt solution (HBSS, Solarbio, Beijing, China) for three times, cells were incubated for 30 min with 2 mL HBSS, which was added with 2μL 1 mg/mL fura-2/AM (Invitrogen, Carlsbad, CA, USA, dissolved in DMSO,) under shaking conditions (60 rpm) at room temperature in dark. Subsequently, fura-2/AM solution was removed and cells were then washed twice with HBSS and were covered with 2 mL of fresh HBSS. Imaging experiments were conducted on a Leica live cell imaging microscope (DMI6000B) equipped with a CCD camera. Cells were stimulated with 340 and 380 nm UV light. A 200 μL test solution was added into the dish using pipette. Data acquisition and analysis were performed by using Leica LAS-AF software (version 2.6.0). The F_340_/F_380_ ratio was analyzed to measure Ca^2+^. Cell assays were repeated for at least three times. Sample size of cell assays to achieve adequate power was chosen on the basis of a previous report with similar methods (Sato *et al.*, 2011).

#### Microscale thermophoresis

In this study, the Microscale Thermophoresis device (Nano Temper Technologies) was used for measurements of binding affinities. Briefly, the recombinant plasmid pIZ-NlGr23a-GFP was constructed using the *NlGr23a* ORF with Kozak consensus sequence at the 5’ ends and *GFP* coding sequence at the 3’ ends cloned into pIZ-V5-His vector. Subsequently, Sf9 cells were transfected with the pIZ-NlGr23a-GFP plasmid (2 μg per 10^6^ cells) using Lipo3000. To achieve higher transfection efficiency, a second transfection was performed in the same batch of Sf9 cells at 24 h after the first transfection. Cells were harvested 48 h after the second transfection. After one wash in PBS, cells were lysed in RIPA lysis buffer and then incubated on a rotary table. After samples were rotated for 1.5 h, the lysates were centrifuged at 13,000 xg for 15 min at 4°C. The extract was harvested and separated into 5 μL per portion; each portion of cell lysate was mixed gently with equal volume of 2 fold gradient diluted OA or glycerol respectively and used for microscale thermophoresis under procedures described previously (Wienken *et al.*, 2010).

### RNAi treatments

#### dsRNA preparation and injection into BPH

Template DNA for dsRNA synthesis was amplified using gene-specific primers (see see Appendix 2—table 1). The resulting purified products were then used to synthesize dsRNA using a MEGAscript T7 High Yield Transcription Kit (Promega, Madison, WI, USA). The concentration of *NlGr23a* dsRNA (239 bp) was quantified with a NanoDrop 2000 instrument (Thermo Fisher Scientific, Waltham, MA, USA). Finally, the quality and size of the dsRNA were further verified via electrophoresis in a 1% agarose gel. *GFP* dsRNA was used as a control. BPHs were collected from the culture chamber and anaesthetized with CO_2_ for 20 s. Approximately 250 ng dsRNA was injected into each individual. After injection, the BPHs were reared on fresh rice plants, which were maintained in transparent polycarbonate jars measuring 15 cm in diameter and 70 cm in height. Individuals exhibiting motor dysfunction and/or paralysis in 12 h were excluded for next steps. Total RNA of whole body was extracted from five randomly selected individuals to examine the gene silencing efficiency at different time-points (24 h, 48 h and 72 h) by quantitative reverse transcription PCR (qRT–PCR). Total RNA of antennae, mouthparts and legs separately dissected from 30 females 48 h after ds*NlGr23a* injection was subjected to examine the gene silencing efficiency in primary gustatory organs. Feeding bioassays were conducted 48 h after injection.

#### qRT-PCR analysis

The qRT-PCR procedure was performed using a Light Cycler 480 (Roche Diagnostics, Basel, Switzerland) with a SYBR®FAST Universal qPCR Kit (KAPA, Woburn, MA, USA), following the manufacturer’s instructions. Each reaction mixture included 1 μL of cDNA template equivalent to 1 ng of total RNA, 0.3 μL of each primer (10 μM), and 5 μL SYBR mix in a total volume of 10 μL. Reactions were performed in triplicate for each sample; three biological replicates and three reactions for each biological replicate were performed. Gene expression levels were normalized to the expression level of BPH *β*-*actin* (Chen *et al.*, 2013). The specific primers used for qRT-PCR are listed in Appendix 2—table 1. The amplification conditions were as follows: 95 °C for 5 min, followed by 45 cycles of 95 °C for 10 s, 60 °C for 20 s and 72 °C for 20 s.

### Behavioral experiments

#### Feeding bioassays of BPHs on artificial diet

The no-choice test apparatus for evaluation of feeding decision behaviour was a BPH rearing chamber and artificial diet D-97 (Figure 2A, top) (Fu *et al.*, 2001). Two layers of stretched parafilm were filled with artificial diet (40 μL), either with OA (100 μM, +OA) or without (-OA). BPHs (n =6-10) were placed onto the food-containing parafilm at the beginning of the experiment in each of three replicates. We counted the insects remaining on each parafilm (N_on_) at 30-min intervals for 4 h. The proportion of BPHs accepting food sources was calculated as (food acceptance = N_on_/N_total_), where N_total_ represents the total number of test insects. In two-choice tests, parafilms containing 40 μL of either +OA or -OA artificial diet were located at opposite open ends of the chamber. Ten BPHs per replicate were introduced into the middle of the cylinder at the beginning of the experiment; three replicates were performed (Figure 2B, top). Insects were starved for 6 h before each experiment. We scored the distribution of insects on each side at 30-min intervals for 4 h. Values were totalled and a position index (PI) for the 4 h period was calculated as (PI = (BPHs on the food with OA - BPHs on the food without OA) / (BPHs on the food with OA + BPHs on the food without OA)). The PI values range from −1 to 1, with PI = 0 indicating no position preference for either control or treatment artificial diets, PI > 0 indicating preference for treatment diets, and PI < 0 indicating preference for control diets. The set-up was placed in a walk-in chamber at 26 ± 2 °C at a relative humidity level of 80% ± 10%.

To directly measure feeding, we performed food choice assays as described previously (Sang *et al.*, 2019; Lee *et al.*, 2010). Briefly, 10 BPHs (1-day-old brachypterous female adults) were starved for 6 h and introduced into a glass cylindrical column in each replicate (8≤n≤10). Then, we prepared two mixtures with each non-toxic dye enclosed in parafilms and located the mixtures at opposite ends of a glass cylindrical column to monitor their food preference. One mixture contained the indicated concentration of OA and 10% (w/v) sucrose with blue dye (Brilliant Blue FCF, 0.2 mg/ml), and the other mixture contained 2% sucrose with red dye (sulforhodamine B, 0.1 mg/ml). BPHs were given 4 h to choose between these two choices. After feeding, BPHs were sacrificed by freezing at −20°C, and then, we dissected them to observe their mid-gut colors under the microscope for the presence of red (N_R_), blue (N_B_), or purple (N_P_) dye. The preference index was calculated using the following equation: preference index = (N_R_ - N_B_)/(N_R_ + N_B_+ N_P_), depending on the dye/tastant combinations. Preference index of −1.0 or 1.0 indicated a complete preference to either 10% sucrose with OA or 2% sucrose alone, respectively. A preference index of 0.0 indicated no bias between the two food alternatives. Each treatment had four biological repeats.

#### Ingestion assay

To assess whether much lower concentration of OA inhibited BPH feeding, the ingestion assay was carried out following a modified protocol (Sang *et al.*, 2019). After 6 h starvation, 1-day-old female BPHs were allowed to feed on 1% (w/v) Brilliant Blue FCF food with or without 100 μM OA for 4 h. BPHs were sacrificed by freezing at −20°C af ter feeding. Six females were transferred into 1.5 ml centrifuge tubes filled with 200 μl PBST (1× PBS with 0.2% Triton X-100) and were completely ground. Then 800 μl of PBST was added and blended. Sample was centrifuged for 5 min at 15,871 g at 4°C. Subse quently, the absorbance of supernatant was measured at 630 nm using a NanoDrop 2000 instrument. PBST alone was used as the blank control. The experiment was repeated with 4 biological replicates.

#### Surgeries

Females were anesthetized with CO_2_, and the second and third antennal segments were removed with a set of sharp forceps.

#### Feeding bioassays of BPHs on rice

Rice with a higher content of OA (TN1^+OA^) was obtained by the immersion method. Briefly, TN1 rice at the tillering stage was immersed in OA solution (20 mM) for 5 min and air-dried for 1 h. The resulting OA level was determined as follows. Firstly, the rice was washed and drained, then ground to a powder using liquid nitrogen. Secondly, the rice powder was placed in a 50 mL centrifuge tube and mixed with 0.5 mol/L HCl (fresh weight of rice : HCl = 1:5; m:m). The homogenate was placed in a boiling water bath for 15–20 min then centrifuged at maximum speed. The resulting supernatant was diluted 10 times with ddH_2_O. Finally, the sample was filtered with a microporous filter membrane (0.45 μm). The OA content was determined using an UltiMate 3000 HPLC (Dionex Corporation, Sunnyvale, CA, USA) equipped with an Agilent ZRBAX SB-C18 column (4.6 mm × 150 mm, 3.5 μm). The mobile phase w as 5 mM TBA in 0.5% KH_2_PO_4_ (pH 2.0) with a flow rate of 1 mL/min. The eluate from the column was continuously monitored at 220 nm after sample loading (5 μL).

To evaluate BPH food preferences across rice diets with different OA levels, we performed dual choice assays as described above with the following modifications. At the beginning of each experiment, ten BPHs were placed on the middle of a sponge, which sealed an upside-down plastic cup containing one TN1 stem and one TN1^+OA^ stem. Probing marks were detected based on a previously described method (Sogawa and Pathak, 1970). In each of four replicates, BPHs (n = 4) reared on one rice stem were confined for 4 h using a 50 mL polypropylene tube. The exposed plant parts were stained with 1% eosin Y (Aladdin Reagent, Shanghai, China), then probing marks on the plant surface were counted.

#### Tarsal avoidance response (TAR) assay

TAR assay referenced proboscis extension response (PER) assay (Lee *et al.*, 2010). After BPH adults were anaesthetized with CO_2,_ the back of each adult was immobilized on a Petri dish using a double-stick tape, while all legs were remained free. BPHs were allowed to fully recover in a humidified incubator for 10-15 min. Before TAR testing, water was given to BPHs to exclude insects in a fright or unhealthy state, if they showed avoidance to water. Then, tastants were delivered to the distal parts of legs without contacting the mouthpart and the antennae using a small piece of Kimwipe paper wicks. TAR response of each individual was observed within 3 s after tatstant treatment. The response, the hindlegs avoiding the contact of Kimwipe, was counted as TAR. BPHs often adhere on wet wicks treated with water or stimulant only and used leg to push away Kimwipe treated with deterrent tastants. Insects were tested in a group of 15–20 individuals, and the percent of insects showing TAR to each tastant was observed and recorded.

To determine the behavioral significance of OA to the tarsal behavior, we contacted the legs of each PBH with 2% sucrose (1^st^ exposure), 2% sucrose plus 5% OA (1^st^ exposure), 2% sucrose (2^nd^ exposure), and 2% sucrose plus 5% OA (2^nd^ exposure), sequentially. TAR response of each individual was observed within 3 s after tatstant treatment. Trials were separated by a 10 min inter trial interval so as to remove any gustatory clues which could have been left behind during previous trials. To determine the deterrent effect of OA, BPHs were treated with different concentrations of OA (10, 20, 50, 100 μM). To test the effect *NlGr23a* on OA detection via the tarsus, ds*NlGr23a*-injected BPHs were applied with 100 μM OA. *GFP* dsRNA was used as a control. Each treatment or control had four biological repeats.

#### Electrical penetration graph (EPG) recording of BPH feeding behaviour

BPH feeding behaviour was recorded on a Giga-8 DC EPG amplifier (GDAAS, Guangdong, China) based on a previously described method (Zhang *et al.*, 2015). Briefly, all experiments were carried out at 26°C ± 1 °C and 70% ± 10% relative humidity under continuous light conditions. The feeding behaviour of individual BPHs on liquid-diet sacs or rice was monitored for at least 3 h. The rice variety or liquid-diet sacs were used alternately, and new insects were used for each recording. Each treatment was replicated 5 times. The signals recorded were analysed using PROBE 3.4 software (Wageningen Agricultural University, Wageningen, The Netherlands). Each feeding behaviour was expressed as the duration of each waveform as a proportion of total monitoring time (%) and the number of times each waveform occurred.

#### Tissue-specificity of *NlGr23a*

Antenna, mouthpart, tarsus, mid-gut, fat body, ovary and epidermis were separately dissected from 20 to 30 one-day old female adults, depending on the size of the organs and tissues. All collected samples were stored immediately in lysis buffer offered by RNA extraction kit and ground with steel beads in a cryogenic grinder. Total RNA was extracted using a Micro Elute Total RNA kit (Omega Bio-tek, Norcross, GA, USA) following the manufacturer’s protocols. RNA quantity was determined as previously descried. Using 100 ng of total RNA, cDNA was synthesized as before for a RT-PCR template. Full length gene-specific primers of *NlGr23a* were used for RT-PCR analysis listed in Appendix 2—table 1. BPH *β*-*actin* gene was used as control. PCR was performed under the following conditions: 94°C for 2 min, 39 cycles of 94°C for 30 s, 55 °C for 30 s, 72°C for 90 s, and 72°C for 5 min. The number of cycles was reduced to 29 for *β*-*actin*. PCR amplification products were run on a 1.2% agarose gel.

#### Statistics

All statistical analyses were performed using IBM SPSS Statistics 24 (IBM, Armonk, NY, USA). The differences between two groups were analysed using two-tailed *t*-test. Differences among multiple groups were analysed using one-way ANOVA followed by Duncan’s multiple range test for multiple comparisons. *P* < 0.05 was marked “*” and *P* < 0.01 was marked “**”. Data were checked for normal distribution using Shapiro-Wilk test.

---

**Figure 1—figure supplement 1.**
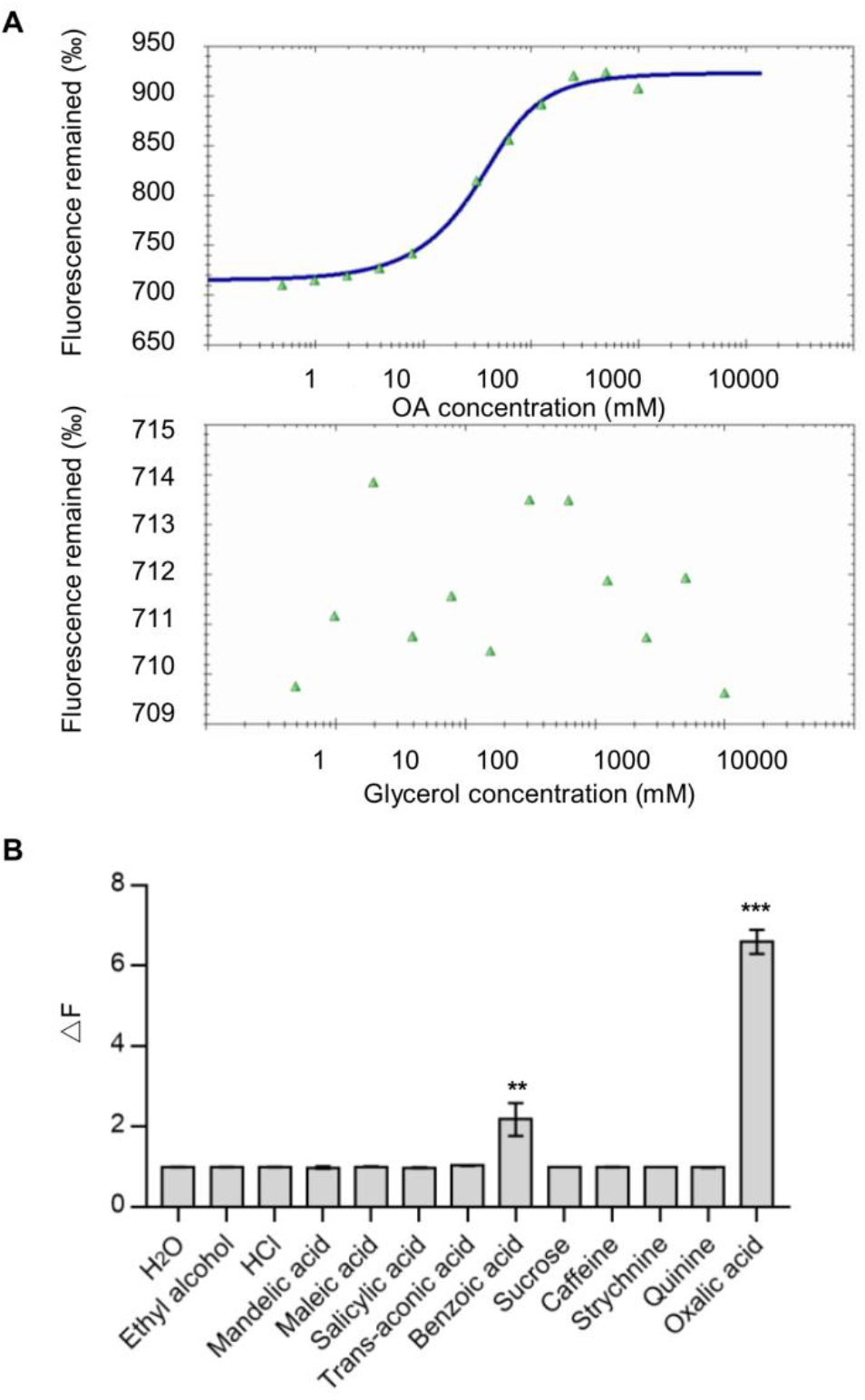
Verification of interaction between NlGr23a and oxalic acid (OA) and response profiles of the HEK293T cells expressing NlGr23a. (A) Verification of interaction between NlGr23a and OA using microscale thermophoresis. Top: The interaction of NlGr23a-GFP with OA. Bottom: The interaction of NlGr23a-GFP with glycerol. Sf9 cells transfected with pIZ-NlGr23a-GFP were harvested and lysed 3 days post transfection. The cell lysates were mixed with equal volume of 2 fold gradient diluted OA or glycerol respectively and used for microscale thermophoresis. (B) Responsiveness of the NlGr23a-HEK293T cells to other phytochemicals. Ca^2+^ response of the NlGr23a-expressing HEK293T cells (n = 50) was stimulated with indicated tastants (each at 10 mM). Bars represent means ± SEM. *t*-test; \*\**P* < 0.01; \*\*\**P* < 0.001.

**Figure 2—figure supplement 1.**
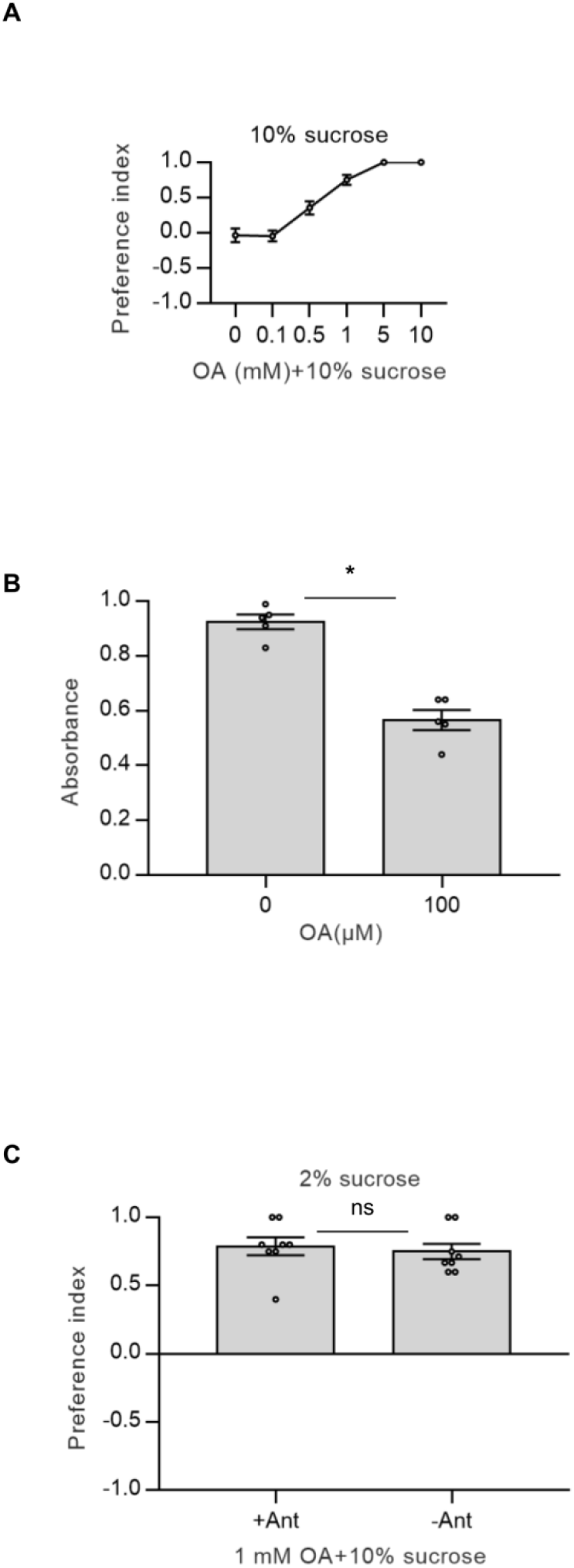
OA induced feeding avoidance. (A) Dose-responsive food choice assay for OA using an equal concentration of sucrose. BPHs were given the choice between 10% sucrose solution and 10% sucrose with different concentrations of OA. The data represent means ± SEM. (n≥8). (B) Effect of OA on food ingestion. Amount of ingestion was calculated using spectrophotogram analysis for diets with or without 100 μM OA. Bars represent means ± SEM. *t*-test; \**P* < 0.05 (n = 5). (C) Role of olfaction in OA avoidance. Bars represent means ± SEM. *t*-test; ns, *P* > 0.05 (n=8).

**Figure 2—figure supplement 2.**
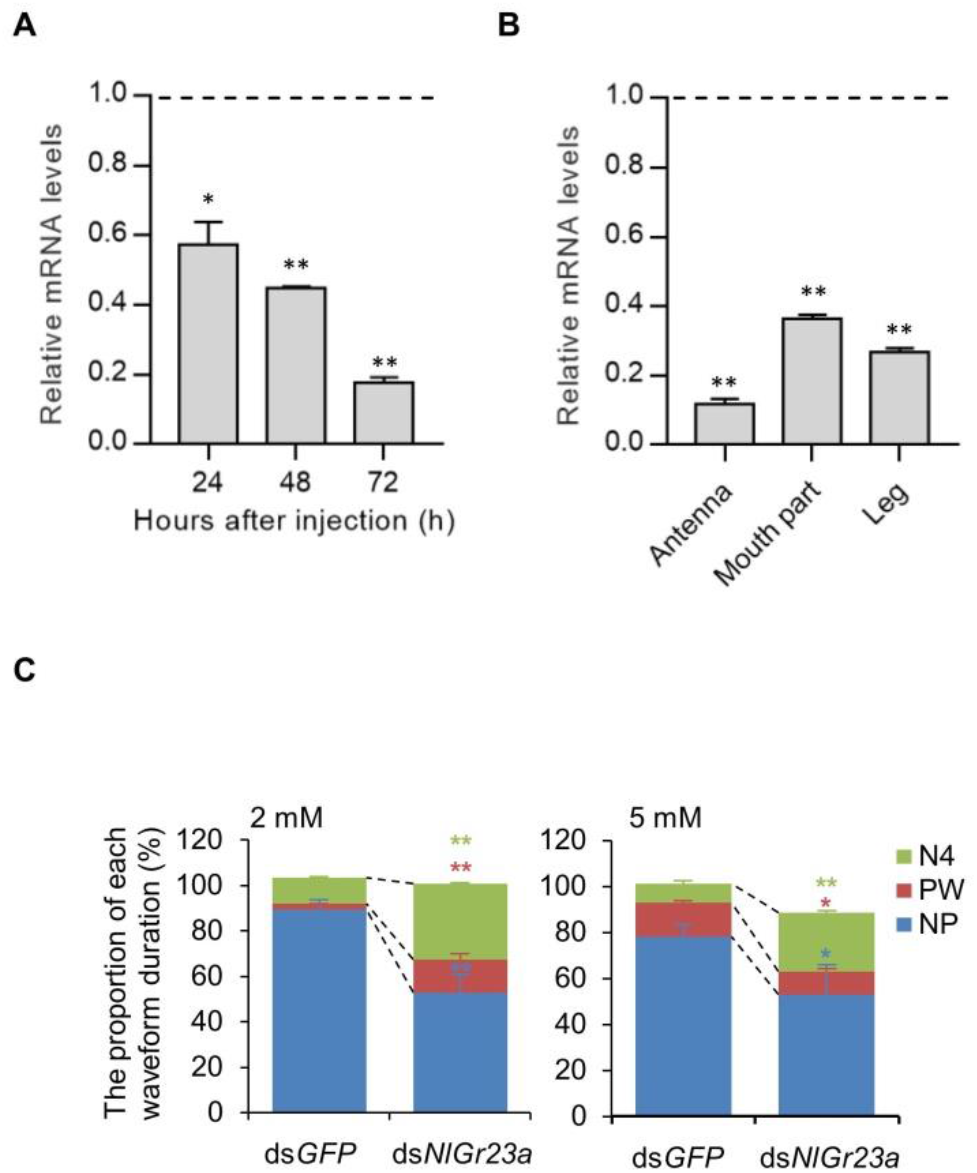
Electrical penetration graph results for BPHs feeding on liquid diet sacs (LDS) after *NlGr23a* silencing and OA content in different rice varieties. (A) The messenger RNA (mRNA) levels of *NlGr23a* in BPH 24, 48 and 72 h after injection of *NlGr23a* dsRNA. All mRNA levels were normalized relative to the *β*-actin mRNA levels. The mRNA level of *GFP* was set to 1. Bars represent means + SEM. *t*-test; \**P* < 0.05; \*\**P* < 0.01 (n = 3). (B) The mRNA levels of *NlGr23a* in primary gustatory organs at 48 h post injection of *NlGr23a* dsRNA. All mRNA levels were normalized relative to the *β*-actin mRNA levels. The mRNA level of *GFP* was set to 1. Bars represent means + SEM. *t*-test; \*\**P* < 0.01 (n =3). (C) The duration of each waveform as a proportion of observation time produced by ds*NlGr23a*- and ds*GFP*-treated BPHs on LDS containing artificial diets with 2 mM or 5 mM OA. Stacked bar graphs represent means + SEM. *t*-test; \**P* < 0.05; \*\**P* < 0.01 (n = 5).

## Appendix 1

**Appendix 1—figure 1.**
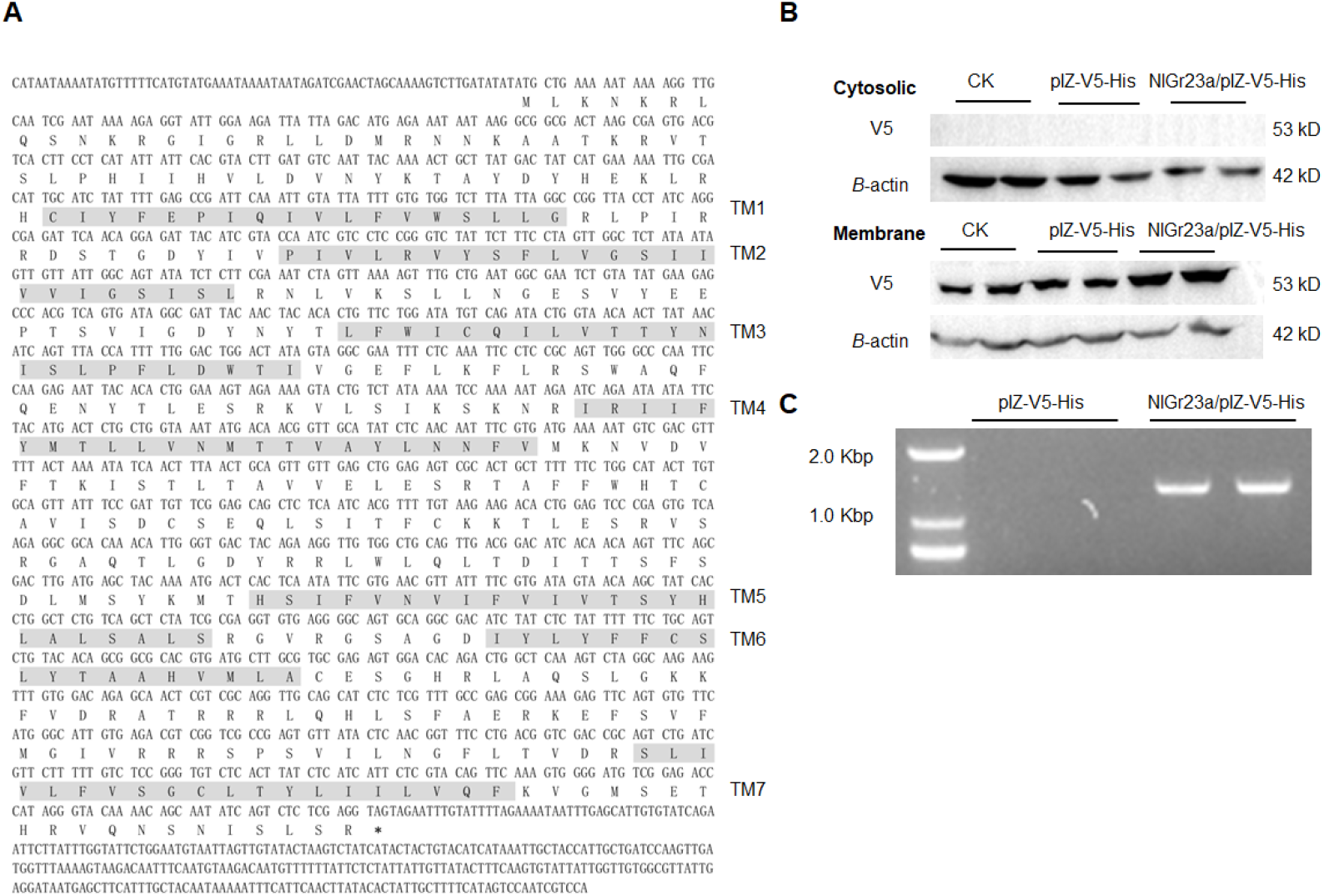
The gene structure of *NlGr23a* and NlGr23a expressed in Sf9 cells. (A) Nucleotide and deduced amino acid sequences of *NlGr23a* cDNA. The predicted seven transmembrane domains (TMs) are shaded in grey. “*” represents the terminal codon. (B) The expression of NlGr23a in the stably transfected Sf9 cell line determined by western blotting. Post antibiotic selection, membrane proteins were isolated (membrane: total membrane proteins; cytosolic: cytosolic proteins). The NlGr23a expression was detected by the anti-V5 antibody. For membrane proteins visualization, the antibody against β-actin was used as the loading control. Stripping protocol was performed using western blot stripping buffer on the same membrane to obtain β-actin signal. NlGr23a/pIZ-V5-His represents pIZ-NlGr23a-V5-His vector transfected cells. CK represents non-transfected cells and pIZ-V5-His represents pIZ-V5-His vector transfected cells as the negative controls. (C) The expression of *NlGr23a* in the stably transfected Sf9 cell line examined by RT-PCR. Post antibiotic selection, RNA from the transfected Sf9 cells was isolated and subjected to RT-PCR analysis of *NlGr23a* transcription.

**Appendix 1— figure 2.**
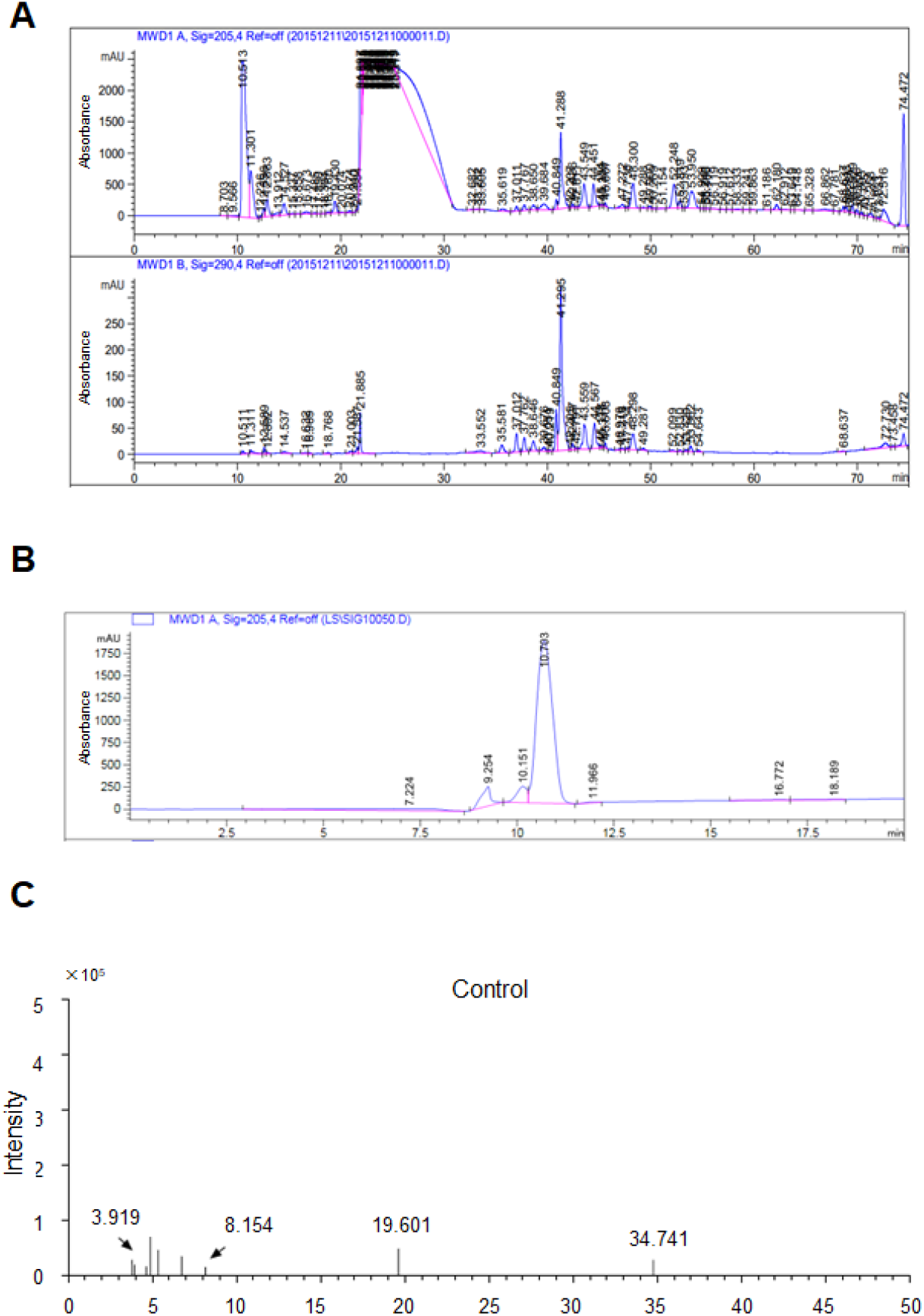
Isolation and identification of rice crude extracts. (A) HPLC chromatogram of the ethyl acetate fraction. The UV detection wavelengths were set at 205 nm (top) and 290 nm (bottom). (B) HPLC chromatogram of bioassay-positive fraction from round 2 of ligand screening. The UV detection wavelength was set at 205 nm. (C) GC-MS chromatogram of the control solution.

**Appendix 1—table 1.** Identification of differential peaks by GC-MS.

| Retention time (min) |  | Intensity |  | Name |
| --- | --- | --- | --- | --- |
| Control | Sample | Control | Sample |  |
| 3.919 | 3.909 | 22228 | 100977 | Oxalic acid |
| 8.154 | 8.144 | 18117 | 445691 | Glycerol |
| 19.601 | 19.581 | 51353 | 62489 | Phthalic acid |
| 34.741 | 34.691 | 31094 | 48765 | Trisiloxane |

## Appendix 2

**Appendix 2—table 1.**
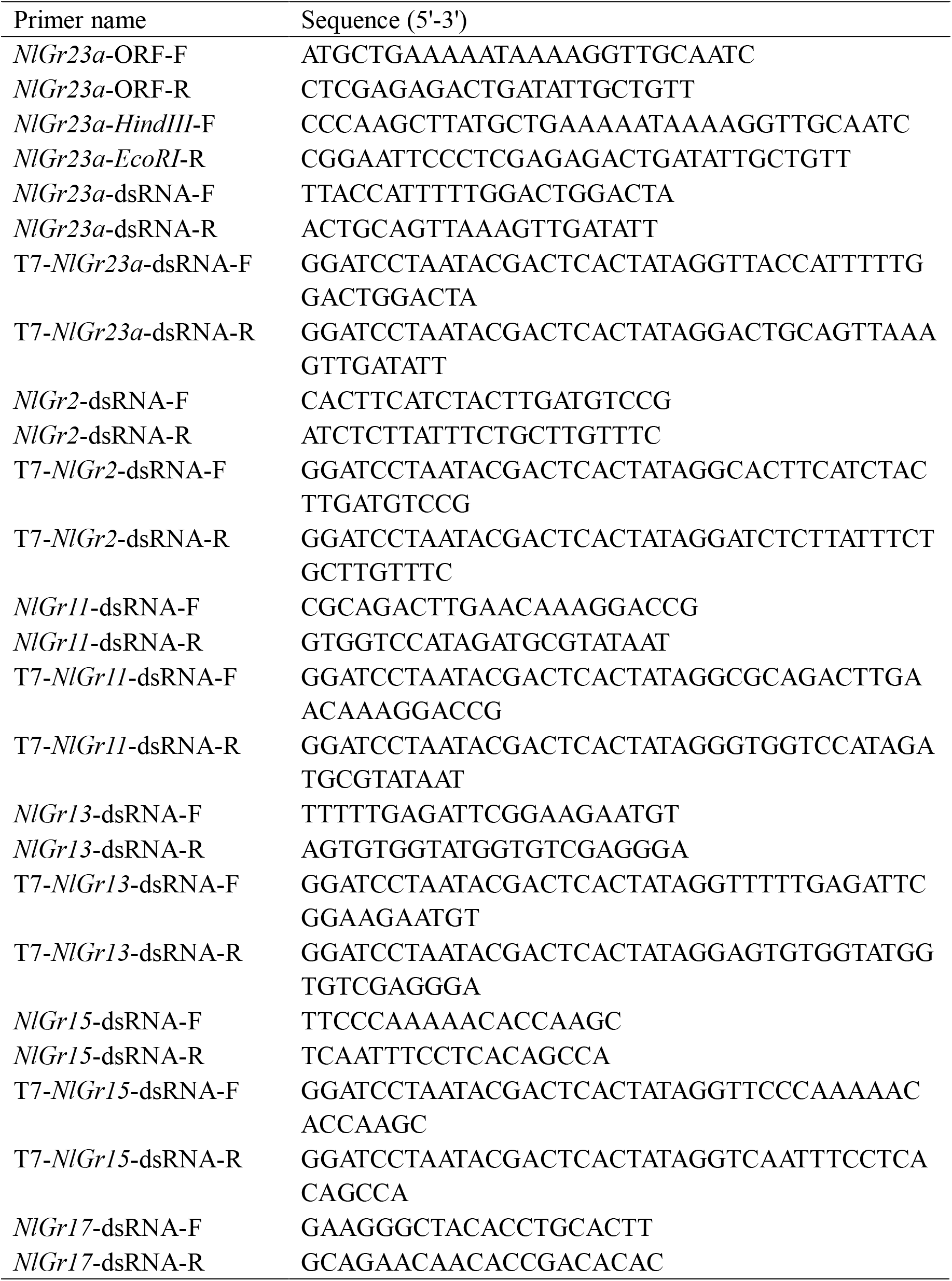
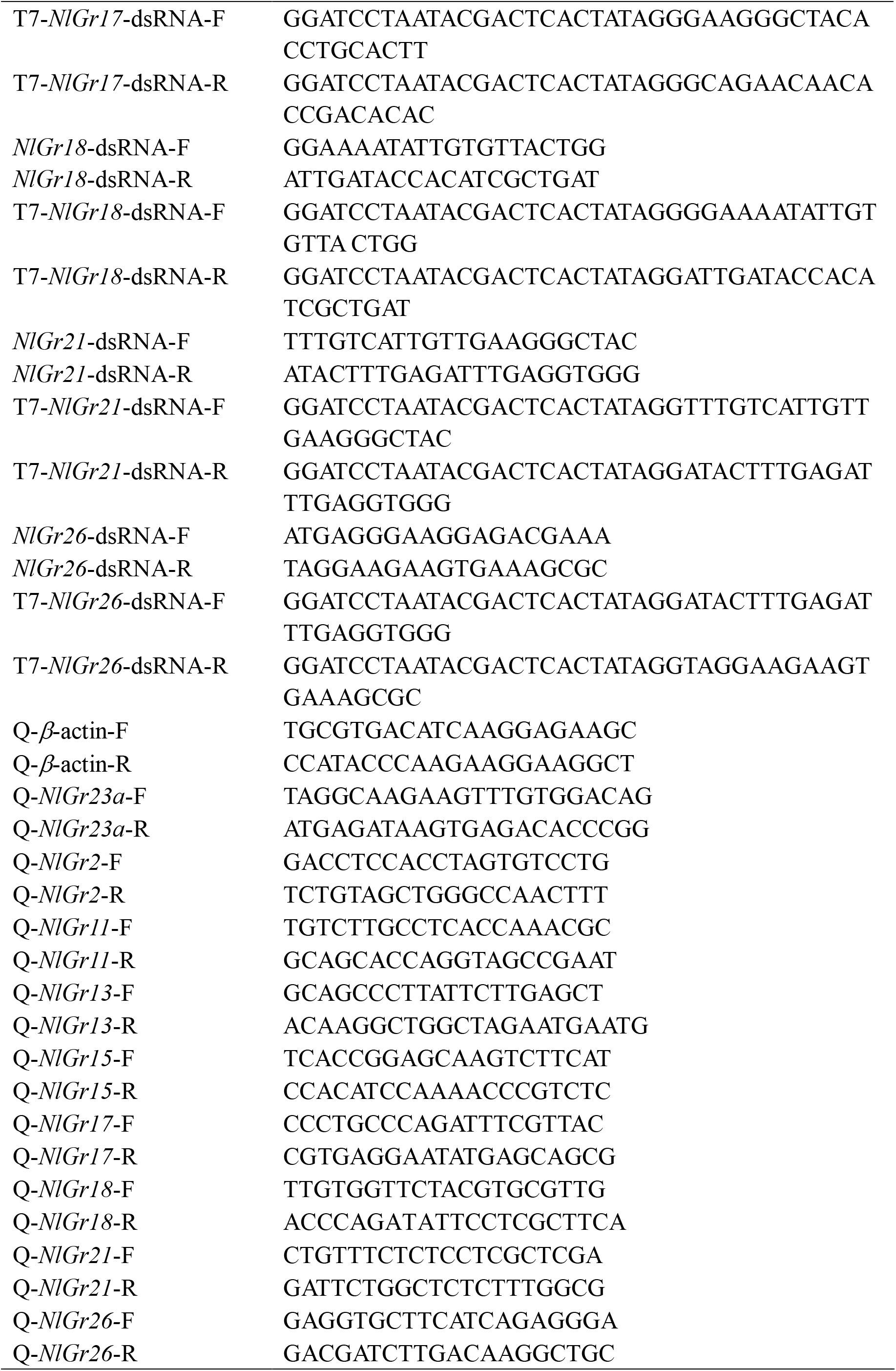
Primers used in this study.

## Appendix 3

**Appendix 3—figure 1.**
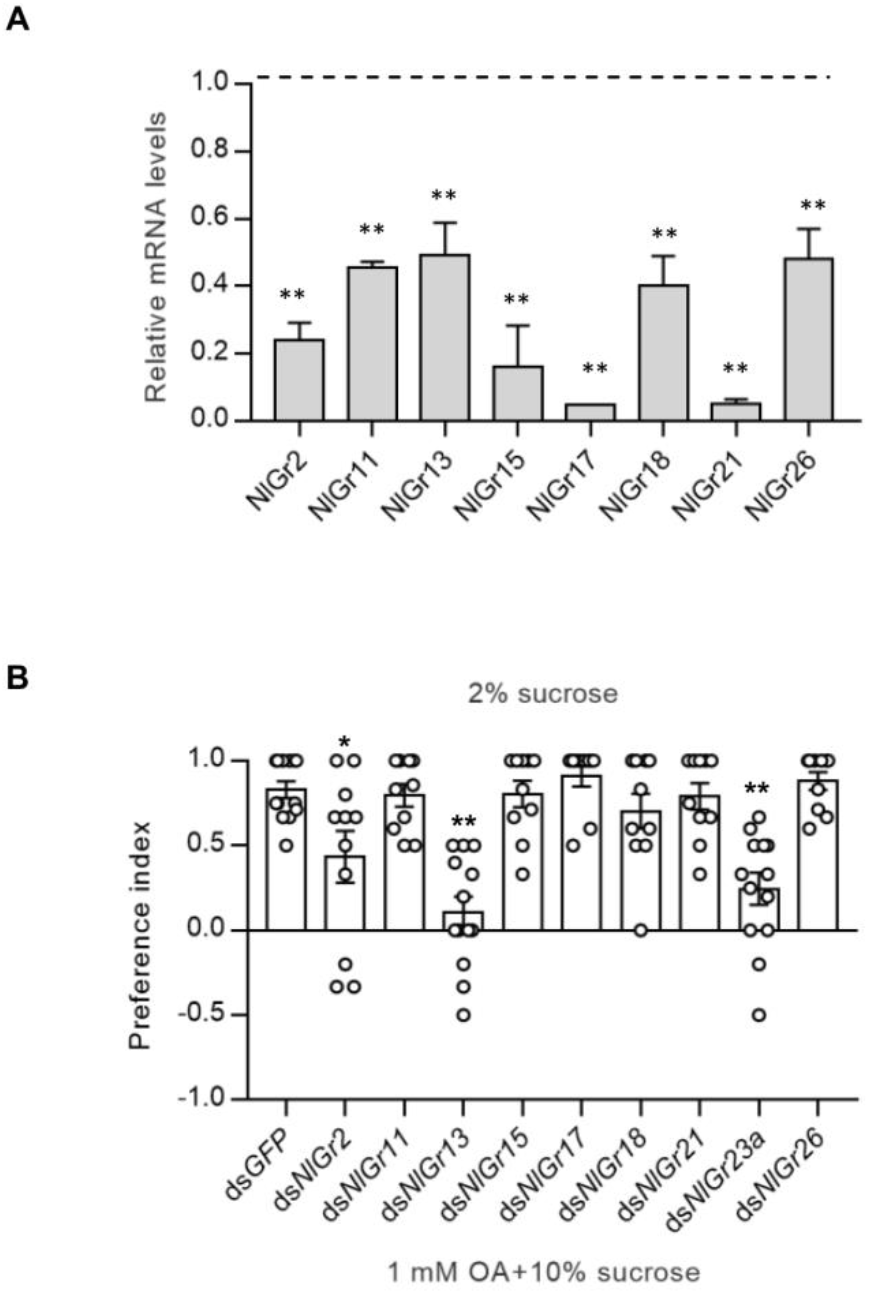
Screen of potential co-receptors of NlGr23a using food choice assays. (A) The mRNA levels of target genes at 48 h post injection of dsRNA. All mRNA levels were normalized relative to the *β*-actin mRNA levels. The mRNA level of *GFP* was set to 1. Bars represent means + SEM. *t*-test; \*\**P* < 0.01 (n =3). (B) Unbiased screen of 8 potential co-recepetors of NlGr23a in binary food choice assays. The data represent means ± SEM. (n≥10). *t*-test; \**P* < 0.05; \*\**P* < 0.01

## Appendix 4

**Appendix 4—figure 1.**
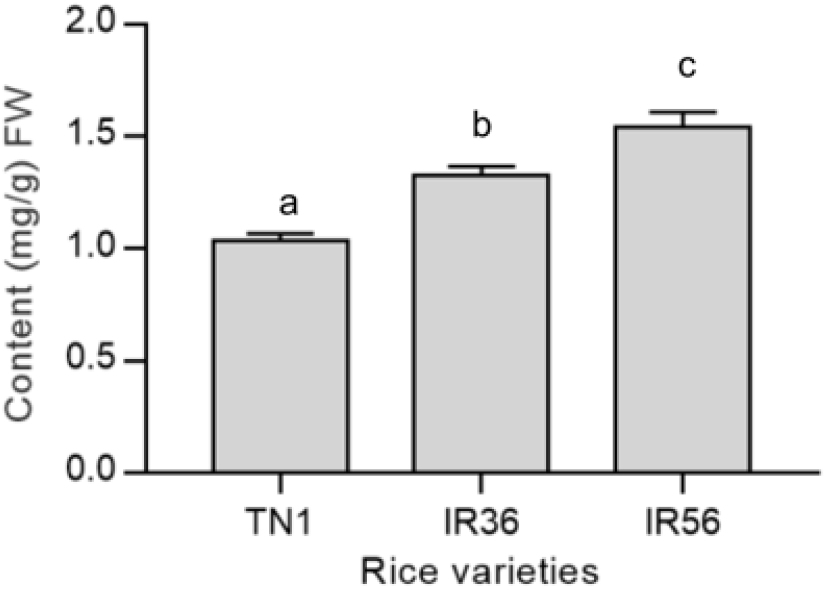
OA content in different rice varieties. OA content in different rice varieties analysed by HPLC. Bars represent means + SEM. Different lowercase letters above columns represent significant differences at *P* < 0.05 from Duncan’s multiple range test (n=4).

## Appendix 5

**Appendix 5—figure 1.**
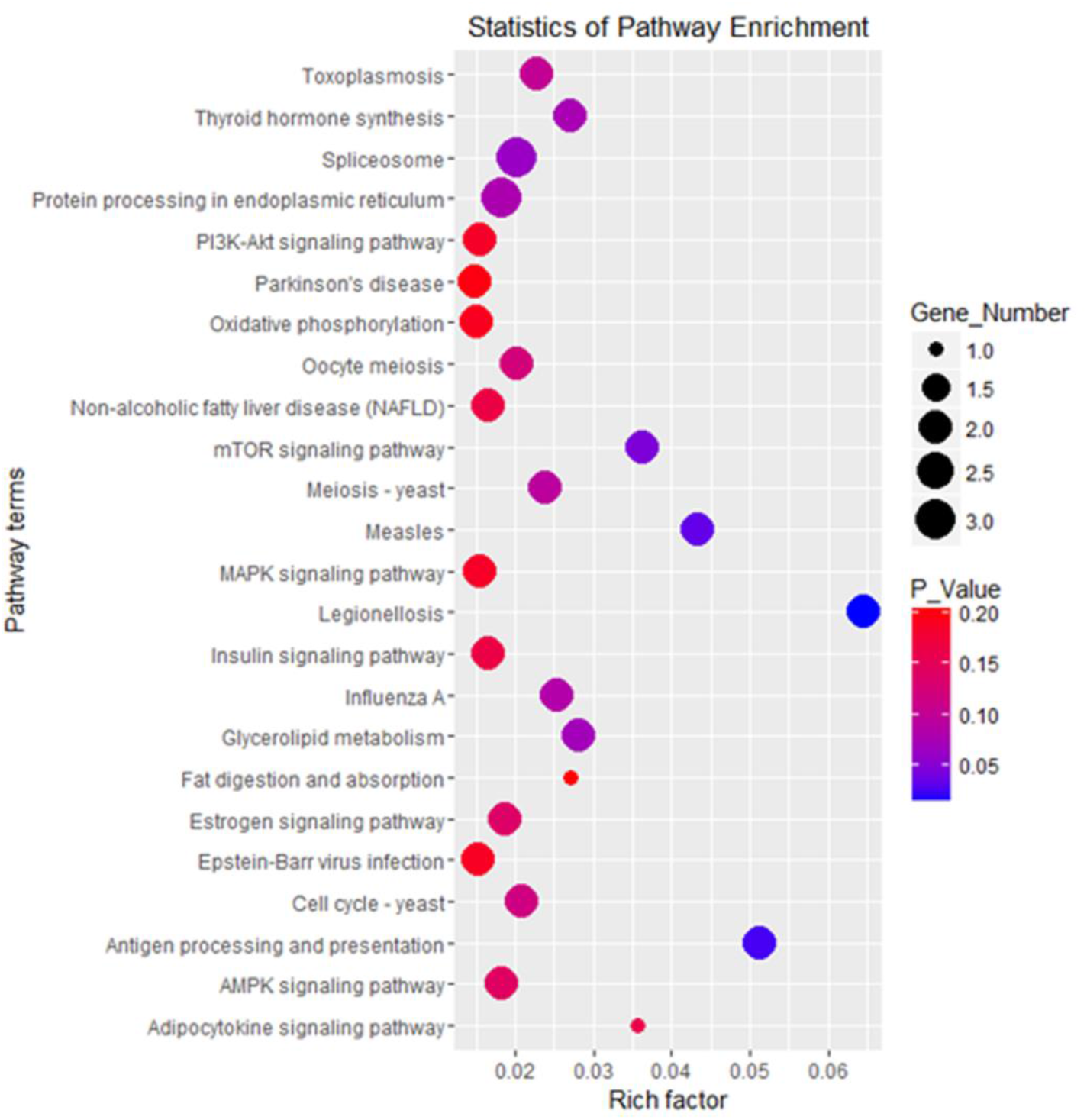
Statistical analysis of enriched pathways for genes differentially expressed between *NlGr23a* dsRNA and *GFP* dsRNA-treated BPHs.

## Appendix 6

**Appendix 6—figure 1.**
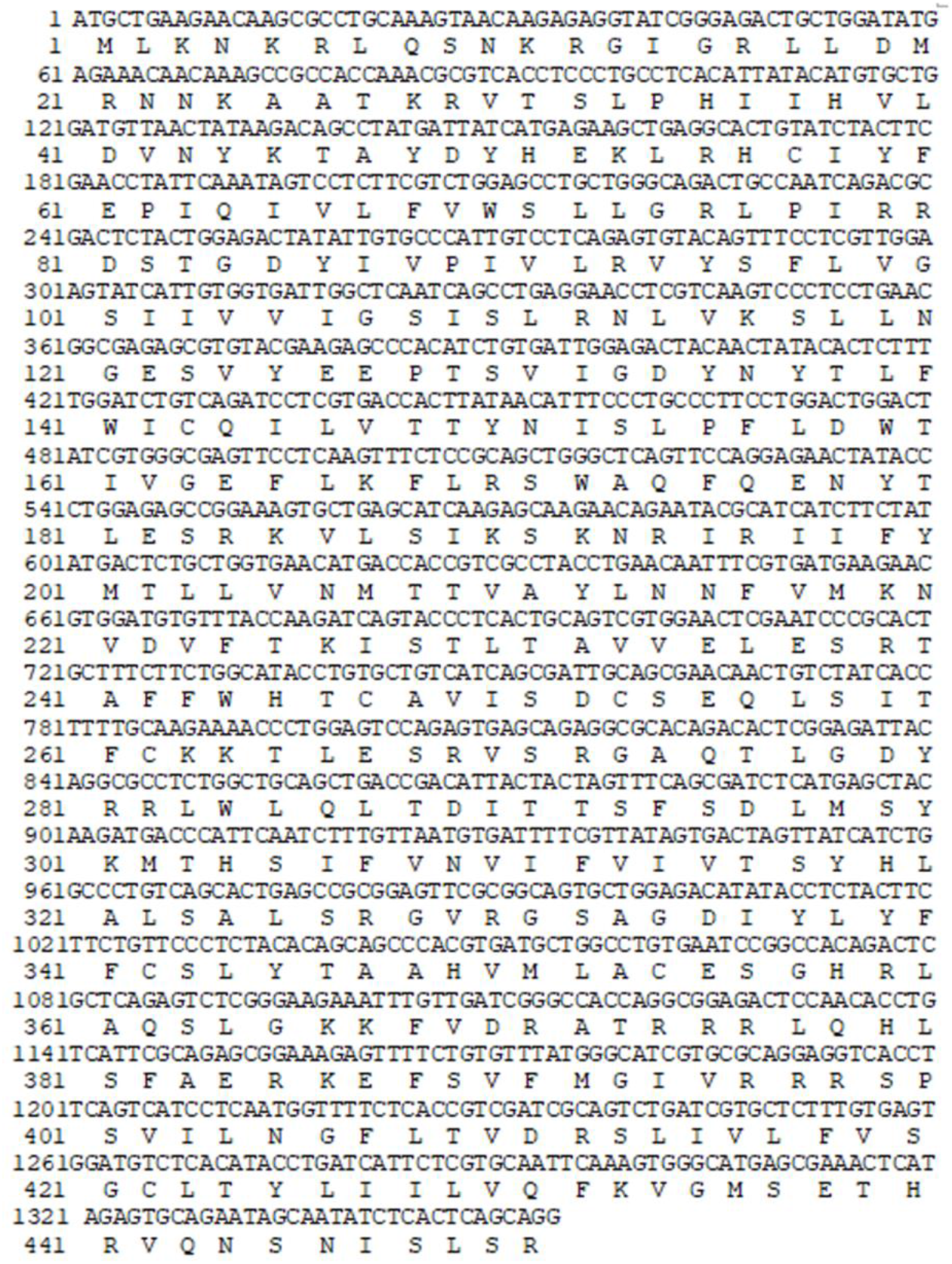
The nucleotide sequences of human codon-optimized *NlGr23a* genes.

## Acknowledgements

We would like to thank Mr. Yang Zhang (Guangdong Academy of Agricultural Sciences, Guangdong, China) for providing us with *N. lugens* strains and his help in using EPG amplifier. This work was supported by the National Natural Science Foundation of China (U1401212, 31672021) and the Young Teacher Foundation of Sun Yat-sen University (19lgpy205).

## Contributions

WZ designed and supervised the entire study. KK carried out ligand identification experiments and feeding bioassays of BPHs on artificial diet. MZ carried out feeding bioassays of BPHs on rice and TAR assays. LY performed the GC-MS and identified compounds. WC helped in developing the calcium-imaging assay method. YD and SX conducted the HPLC separation of the bioassay-active fractions. K. Lin performed and analysed EPG recordings. K. Liu and JL carried out the quantitative RT-PCR. ZG carried out microscale thermophoresis assay. KK and MZ analysed the data. WQ, MZ and KK wrote the manuscript. All authors read and commented on the manuscript.

